# Breathlessness catastrophising after COVID-19 involves both interoceptive and visceromotor connectivity

**DOI:** 10.64898/2026.09.11.750997

**Authors:** Lin Qiu, Betty Raman, Mark Cassar, Emily Fraser, Lauren Z Atkinson, Stuart Clare, William T Clarke, Stefan Neubauer, Helen E Davies, Neil A Harrison, C John Evans, Martyn Ezra, Kyle TS Pattinson

**Affiliations:** Nuffield Department of Clinical Neurosciences, University of Oxford, Oxford, United Kingdom; Nuffield Division of Anaesthetics, University of Oxford, Oxford, United Kingdom; Oxford Centre for Integrative Neuroimaging (OxCIN), FMRIB, Nuffield Department of Clinical Neurosciences, University of Oxford, Oxford, United Kingdom; Radcliffe Department of Medicine, University of Oxford, Oxford, United Kingdom; Oxford University Hospitals NHS Foundation Trust, Oxford, United Kingdom; Department of Psychiatry, University of Oxford, Oxford, United Kingdom; Cardiff and Vale University Health Board, Cardiff, United Kingdom; Cardiff University Brain Research Imaging Centre (CUBRIC), Cardiff University, Cardiff, United Kingdom

## Abstract

Persistent breathlessness is common after COVID-19, yet its severity typically correlates weakly with objective clinical measures. This discordance is often attributed to perceptual inference amplified by anxiety. Because breathing involves a closed perception–action loop, breathlessness may also reflect alterations in interoceptive and visceromotor pathways. We acquired 7-tesla resting-state functional MRI in 53 post-COVID patients with varying breathlessness and estimated functional connectivity across 18 pre-defined interoceptive and visceromotor regions. Linear regression identified two connections associated with breathlessness catastrophising: reduced dorsal periaqueductal grey–posterior insula (dPAG–PoI1) and increased basolateral amygdala– dorsal anterior cingulate (BLA–dACC) connectivity. The dPAG–PoI1 association was stronger in patients who had required mechanical ventilation, whereas the BLA–dACC association was attenuated at higher generalised anxiety. These findings are consistent with two potentially separable contributions to symptom burden: weakened interoceptive signalling between brainstem and sensory cortex, and heightened visceromotor influence of threat processing on autonomic control, rather than anxiety-driven misperception alone.

## Introduction

### Breathlessness uncoupled from pulmonary function

Breathlessness (dyspnoea) is a distressing and disabling symptom that cuts across a wide range of clinical presentations, from chronic respiratory disease to cardiometabolic and functional syndromes (Gysels and Higginson, 2008; Hutchinson et al., 2018; Williams and Carel, 2018). Crucially, its severity is not always coupled to objective clinical measures of pulmonary impairment: patients can experience marked breathlessness despite preserved spirometry and minimal structural lung pathology (Han et al., 2004; Hull and Haines, 2022). This maladaptive symptom–biomarker discordance is clinically consequential. Dyspnoea is a strong predictor of reduced quality of life (Burgel et al., 2013), a more reliable indicator of adverse prognosis in cardiorespiratory illnesses than objective physiological markers such as forced expiratory volume in one second (FEV₁) in COPD (Nishimura et al., 2002; Abidov et al., 2005), and predicts all-cause mortality in the general population in a severity-dependent manner (Figarska et al., 2012; Frostad et al., 2006), yet routine physiological tests provide limited explanatory value for the individual patient. When symptoms outrun what the lungs can explain, the mechanisms maintaining breathlessness are likely to lie, at least in part, in how respiratory signals are centrally processed and regulated, and identifying these neurobiological mechanisms remains a major challenge.

Persistent breathlessness following COVID-19 offers a tractable setting in which we can examine this discordance. It has emerged as a major clinical problem across the full spectrum of acute illness severity, including those with mild infection who did not require hospitalisation (Ayoubkhani et al., 2022; Sudre et al., 2021; Townsend et al., 2021; Seeßle et al., 2022). Conventional pulmonary investigations, and state-of-the-art assessments of alveolar–capillary membrane function using hyperpolarised xenon MRI, have not consistently identified abnormalities that account for the symptom (Ng et al., 2026; Stewart et al., 2023; Matheson et al., 2022). Where subtle abnormalities are detected, they do not reliably distinguish patients who remain breathless from those who do not, and longitudinal improvement in cardiopulmonary measures does not associate with improvement in symptoms (Cassar et al., 2021; Ng et al., 2026).

Peripheral impairment alone therefore appears insufficient to explain why some individuals remain markedly breathless. Psychological comorbidities are common: 25–49% of individuals experiencing post-COVID breathlessness present with mood disorders or anxiety (Evans et al., 2022; Seeßle et al., 2022), and they are at increased risk of developing such conditions (Taquet et al., 2021; Townsend et al., 2021). This combination of persistent respiratory symptoms, limited explanatory power of pulmonary abnormalities, and frequent affective comorbidity makes post-COVID breathlessness a well-suited setting in which to ask how central regulatory networks contribute to symptom experience.

### Breathing as a closed-loop interoceptive–allostatic process

A dominant account of maladaptive breathlessness emphasises sensory–affective amplification, whereby anxiety symptom severity and negative affect increase the salience of breathlessness sensations and promote self-reinforcing cycles of symptom escalation (Bailey, 2004; Evans, 2010; Strang et al., 2014; Banzett et al., 2020). More recent work has framed these ideas within predictive coding or Bayesian models, proposing that exaggerated breathlessness can arise when catastrophic expectations or threat appraisals outweigh interoceptive evidence, potentially through altered precision-weighting of priors and afferent signals (Harrison et al., 2021; Van den Bergh et al., 2017; Faull et al., 2017). Within these traditions, breathlessness is treated as a problem of perceptual inference, much like vision or other extensively studied sensory modalities.

Breathing, however, is unique among perceptual experiences in that it forms a closed perception– action loop. The afferent signals the brain reads are largely the product of the motor act it has just performed, and each act is shaped in turn by that signal, from the moment-to-moment adjustment of a single breath to slower behavioural strategies such as pacing, overbreathing, or avoidance of movement. Perception and action are thus coupled across timescales, from the respiratory cycle itself to sustained patterns of respiratory behaviour. An account that considers respiratory perception in isolation from respiratory action therefore captures only part of the system and is unlikely to fully explain a symptom that arises when the coupling between the two is disturbed. Characterising the neural basis of maladaptive breathlessness accordingly requires attention not only to regions that sample interoceptive respiratory signals, but also to those that generate the visceromotor predictions and actions with which those signals are integrated.

Interoceptive–allostatic frameworks offer such an integrative account, treating respiratory perception and control as components of a single closed-loop regulatory system (Barrett and Simmons, 2015; Kleckner et al., 2017; Seth and Friston, 2016; Pezzulo et al., 2018). Within these models, interoceptive representations of respiratory state, including signals related to CO₂, pH, and mechanical stretch, are reciprocally coupled with visceromotor control policies: predictions about bodily needs and threat guide respiratory regulation, while afferent signals continuously update those predictions to maintain physiological stability. Critically, this functional division may be grounded in the laminar organisation of the underlying cortex. Kleckner and colleagues (2017) proposed that granular and dysgranular regions, including the posterior and dorsal insula, support the interoceptive sampling of bodily signals, whereas agranular regions, including the anterior cingulate and anterior insula, generate visceromotor predictions and regulatory actions. This laminar distinction provides an anatomical basis on which the two arms of the loop can be identified and empirically tested.

### The present study

Building on this framework, we assembled a set of regions of interest spanning both the interoceptive and visceromotor sides of this loop in breathing perception and regulation. It comprises the primary interoceptive and visceromotor cortical regions characterised above, extended to higher-order and primary sensorimotor cortices, limbic circuitry involved in threat learning and valuation (LeDoux, 2007; Phelps and LeDoux, 2005), and midbrain structures such as the periaqueductal grey subdivisions that integrate defensive and respiratory control (Faull et al., 2019; Laviolette and Laveneziana, 2014). Altered interactions within this network may contribute to persistent breathlessness in the absence of major pulmonary dysfunction.

Resolving these interactions requires sensitive measurement of connectivity within small nuclei and fine-grained insular subdivisions that are difficult to resolve at conventional field strengths. Ultra-high-field 7-tesla resting-state fMRI, particularly with extended acquisitions, offers the improved signal-to-noise ratio and spatial specificity needed to characterise small nuclei, including PAG subregions (Faull et al., 2016; Faull and Pattinson, 2017), and insular organisation central to interoceptive–allostatic regulation. Whereas much of the neuroimaging literature on breathlessness has characterised phasic responses to experimentally induced respiratory challenge, largely in healthy participants, in whom the evoked sensation is a proportionate and adaptive response to an acute stimulus (Faull et al., 2016; Faull and Pattinson, 2017; Herigstad et al., 2011), resting-state imaging allows us to examine intrinsic network organisation in the ongoing symptomatic state, in which breathlessness persists without an identifiable stimulus and takes a tonic rather than phasic form.

In this study, we recruited a cohort of patients with a range of acute COVID-19 severity, comprising both hospitalised and non-hospitalised individuals, and tested whether breathlessness was associated with resting-state functional connectivity among these pre-defined regions. We distinguished threat-related appraisal of respiratory sensations from the reported experience of breathlessness itself and asked whether connections tied to generalised anxiety could be distinguished from those tied to the severity of the acute infection.

## Results

Fifty-three post-COVID patients completed the study (mean age 50 ± 14 years; 59% female; mean body mass index 28 ± 5 kg/m²). During acute infection, 36% (n=19) had been hospitalised, of whom 74% (n=14) had required mechanical ventilation, and the mean WHO Clinical Progression Scale (Marshall et al., 2020) score was 2.4 ± 2.0. Spirometry was available for 33 of the 53 participants (Oxford hospitalised, 11 of 19; Oxford non-hospitalised, 1 of 9; Cardiff non-hospitalised, 21 of 25). In these participants, values were within normal limits (mean FEV₁ 95%, FVC 104%, FEV₁/FVC 79%; Appendix 1). Assessments were conducted a median of 257 days after infection (IQR 179–371; range 125–806; mean 323 ± 196), available for 51 of the 53 participants. Corrected sample sizes were used in the relevant sensitivity analyses (see Methods). Full participant demographics are summarised in Appendix 2.

### Edge-wise associations with breathlessness catastrophising and Dyspnoea-12

We used robust (Huber M-estimator) linear regression to test whether functional connectivity within the interoceptive–allostatic breathing network was associated with post-COVID breathlessness. The analysis included 18 regions of interest (Appendix 5), yielding 153 pairwise connections. Breathlessness was indexed using the Breathlessness Catastrophising Scale and the Dyspnoea-12. All models controlled for age, sex, body mass index, absolute and relative head motion, and cube-root-transformed intracranial and total brain volumes. P-values were corrected for multiple comparisons using the Benjamini–Hochberg false discovery rate (FDR) procedure (Benjamini and Hochberg, 1995) across all 153 tests within each outcome; BCS and D-12 were treated as pre-specified separate primary outcomes and were not corrected across.

For the Breathlessness Catastrophising Scale (BCS; Solomon et al., 2015), regression coefficients for all pairwise connections are shown in Figure 1. Two connections survived FDR correction (p < .05). Specifically, BCS scores were negatively associated with connectivity between the dorsal periaqueductal grey (dPAG; Faull et al., 2016) and posterior insula subdivision 1 (PoI1, a ventral-posterior subdivision of the posterior insula defined in the Glasser atlas; Glasser et al., 2016) and positively associated with connectivity between the basolateral amygdala and dorsal anterior cingulate cortex (BLA–dACC; Figure 1). For the Dyspnoea-12 (D-12; Yorke et al., 2010), no edge survived FDR correction (Figure 2). Accordingly, the subsequent edge-specific analyses focus on these two BCS-associated edges and should not be interpreted as evidence for connectivity correlates of global dyspnoea severity.

Despite this difference in statistical significance, the overall pattern of associations was broadly similar across the two breathlessness measures. Negative associations were predominantly observed between interoceptive regions, including dPAG–PoI1 and the dorsal periaqueductal grey and lateral ventral anterior insula (dPAG–lvaIns). In contrast, positive associations were primarily observed among visceromotor regions, including BLA–dACC and the ventrolateral periaqueductal grey and subgenual anterior cingulate cortex (vlPAG–sgACC), as well as within insular regions such as the dorsal posterior and dorsal medial insula (dpIns–dmIns).

**Figure 1.**
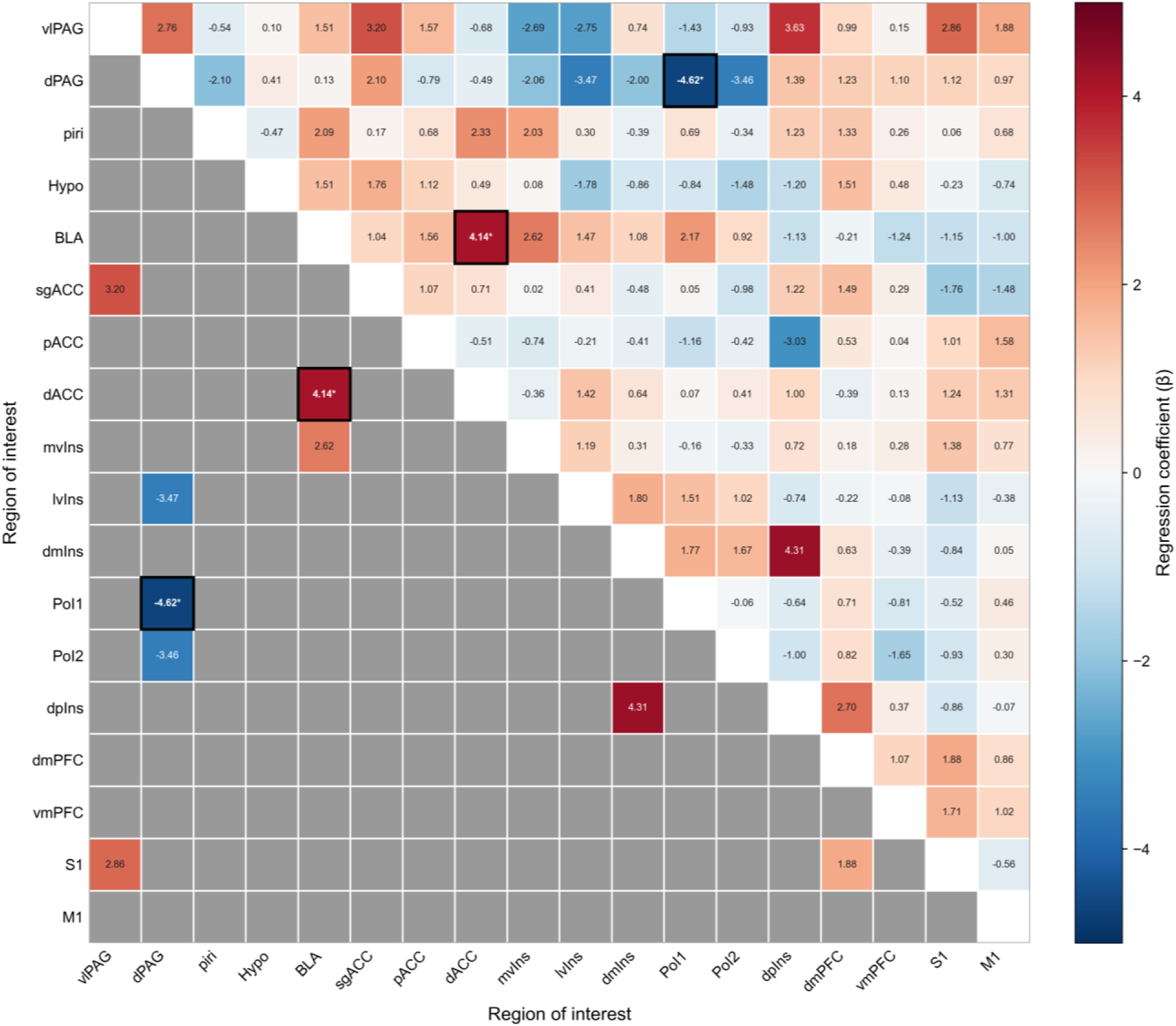
Breathlessness-catastrophising regression coefficients across functional connectivity edges. Each cell represents the coefficient from a univariate robust regression of breathlessness catastrophising scale (BCS) scores on resting-state functional connectivity between the two regions of interest (ROIs) indicated on the x- and y-axes. The upper triangle displays all regression coefficients, whereas the lower triangle displays only coefficients with an unadjusted *p* < 0.05. ROIs are ordered by anatomical region (subcortical, ACC, insula, and cortical regions). The two edges surviving false discovery rate (FDR) correction, dPAG–PoI1 and BLA–dACC, are marked with an asterisk. ROI abbreviations are defined in Appendix 5.

**Figure 2.**
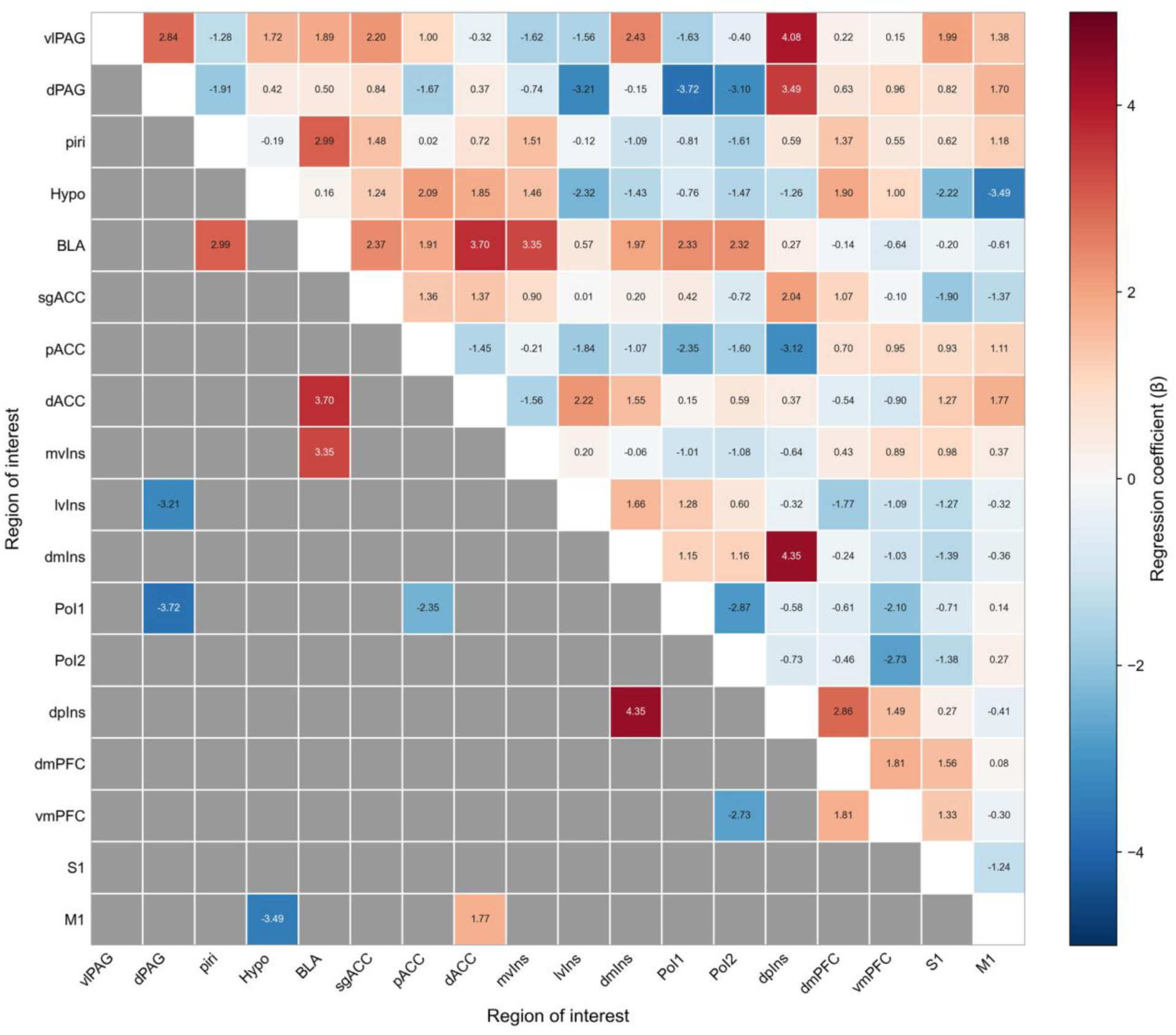
Dyspnoea-12 regression coefficients across functional-connectivity edges. Each cell represents the coefficient from a univariate robust regression of Dyspnoea-12 (D-12) scores on resting-state functional connectivity between the two regions of interest (ROIs) indicated on the x- and y-axes. The upper triangle displays all regression coefficients, whereas the lower triangle displays only coefficients with an unadjusted *p* < 0.05. ROIs are ordered by anatomical region (subcortical, ACC, insula, and cortical regions). No edge survived false discovery rate (FDR) correction. ROI abbreviations are defined in Appendix 5.

### Reduced dPAG–PoI1 connectivity is associated with greater breathlessness catastrophising

Connectivity between the dorsal periaqueductal grey (dPAG) and posterior insula subdivision 1 (PoI1) was negatively associated with BCS (β = −4.62, 95% CI [−6.88, −2.36], p = 6.2 × 10^−5^). We examined the influence of individual observations on this selected edge using leave-one-out (LOO) diagnostics and case-resampling bootstrap confidence intervals (5,000 resamples, bias-corrected and accelerated [BCa] method, seed = 42). The LOO estimate ranged from −5.30 to −3.99 across the 53 folds and did not reverse sign in any fold, and the BCa 95% CI excluded zero ([−8.26, −1.18], p = .036). Nine participants exceeded the conventional DFBETAS threshold (|2/√N| = 0.275); the most influential observation acted to attenuate rather than inflate the association, such that its removal strengthened the effect. These analyses quantify internal stability within the present sample and do not constitute independent, out-of-sample validation.

**Figure 3.**
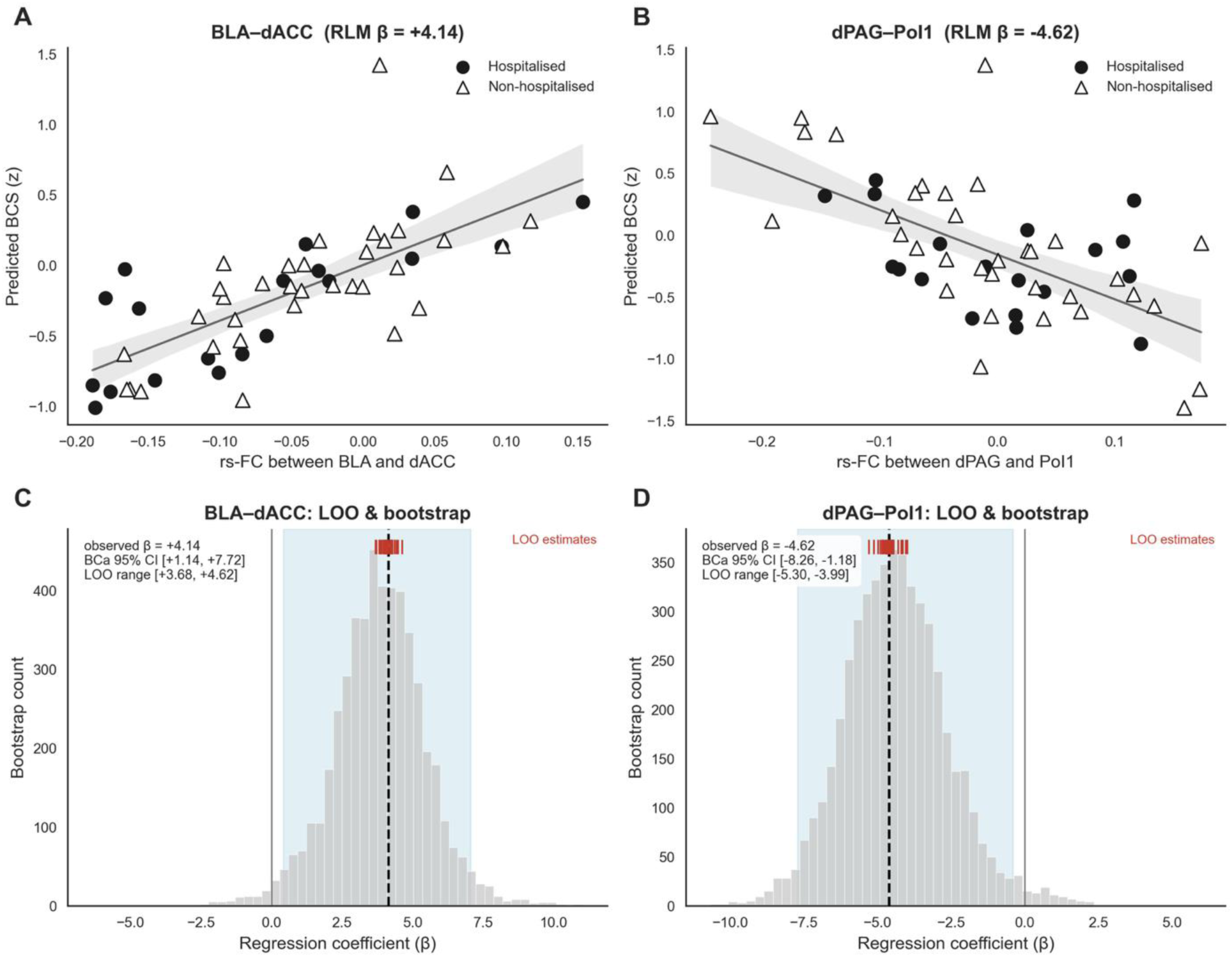
Associations between functional connectivity and the Breathlessness Catastrophising Scale (BCS). (A, B) Scatterplots of resting-state functional connectivity against model-predicted BCS for the two edges that survived FDR correction: BLA–dACC (A) and dPAG–PoI1 (B). Predicted BCS is the fitted value from the primary robust linear regression (Huber M-estimator) of BCS (z-scored) on each edge plus the full covariate set (age, sex, mean absolute and relative framewise displacement, BMI, brain volume, intracranial volume). Filled circles, hospitalised participants; open triangles, non-hospitalised participants; solid line, fitted slope; shaded band, 95% confidence interval. (C, D) Internal-stability diagnostics for the same edges (BLA–dACC, C; dPAG–PoI1, D). Histograms show the case-resampling bootstrap distribution of the regression coefficient (N = 5,000 resamples); the dashed vertical line marks the observed full-sample coefficient and the shaded band the bootstrap 95% percentile interval. Red ticks above each distribution show the 53 leave-one-out (LOO) coefficient estimates, each omitting one participant. Annotations report the observed β, the bias-corrected and accelerated (BCa) bootstrap 95% CI, and the LOO range.

We then examined, as secondary and exploratory analyses, whether the dPAG–PoI1–BCS association was moderated by generalised anxiety scores (GAD-7) or by acute COVID-19 severity (operationalised as requirement for mechanical ventilation). Each moderator was entered as an interaction term with the edge in a robust linear regression (Huber M-estimator), retaining the primary covariate set. Inference on the interaction used non-parametric permutation testing (N = 5,000), with case-resampling bootstrap confidence intervals for the simple slopes.

#### Anxiety

There was little evidence that generalised anxiety scores moderated the dPAG–PoI1– BCS relationship (interaction β = −0.15, p = .88; permutation p = .92; Figure 4, top row).

#### Mechanical ventilation

Mechanical ventilation moderated the dPAG–PoI1–BCS association, with a stronger negative relationship in participants who required ventilation (N = 14; interaction β = −7.24, p = .005; permutation p = .020; Figure 4, bottom row).

**Figure 4.**
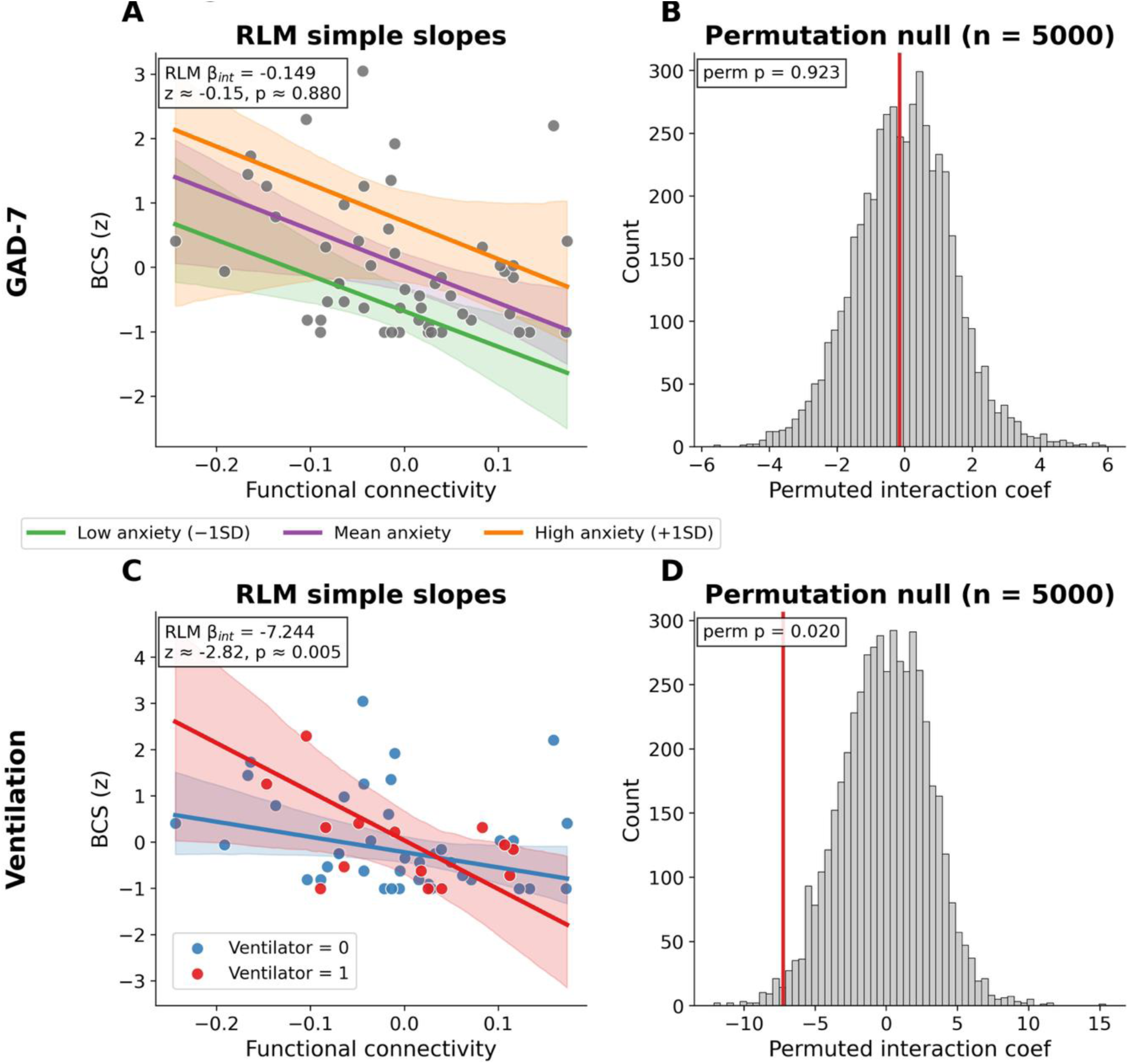
Moderation of the dPAG–PoI1–BCS association. Each moderator was entered as an interaction term with the edge in a robust linear regression model (RLM; Huber M-estimator) retaining the primary covariate set. Top row, generalised anxiety symptoms (GAD-7); bottom row, prior mechanical ventilation. (A, C) RLM simple slopes across the observed range of dPAG–PoI1 connectivity at representative moderator levels (GAD-7: −1 SD / mean / +1 SD, A; ventilated vs non-ventilated, C); shaded bands, 95% case-resampling bootstrap CIs (2,000 resamples); points, individual participants. (B, D) Permutation null distributions (N = 5,000) of the connectivity × moderator interaction coefficient. The vertical red line marks the observed coefficient.

### Increased BLA–dACC connectivity is associated with greater breathlessness catastrophising

Connectivity between the basolateral amygdala (BLA) and dorsal anterior cingulate cortex (dACC) was positively associated with BCS (β = 4.14, 95% CI [1.81, 6.47], p = 5.1 × 10^−4^). Applying the same internal-stability diagnostics, the LOO estimate ranged from +3.68 to +4.62 across folds without sign reversal, and the BCa 95% CI excluded zero ([+1.14, +7.72], p = .031). Three participants exceeded the DFBETAS threshold, and the most influential again acted to attenuate the effect. As above, these diagnostics index internal stability only and are not out-of-sample validation. Moderation analyses used the same procedure as above.

#### Anxiety

Generalised anxiety scores -(GAD-7) significantly moderated the BLA–dACC–BCS association (interaction β = −3.77, p = .020; permutation p = .007; Figure 5, top row), such that the positive association was strongest at lower anxiety and attenuated at higher anxiety scores. Because GAD-7 scores were right-skewed, with half the sample in the minimal range and the higher-anxiety observations clustered in a small severe subgroup (Appendix 7), the simple slope at higher anxiety (evaluated at +1 SD) is estimated across a relatively under-populated region of the moderator. Hence, this moderation should be read as indicative rather than precise.

**Figure 5.**
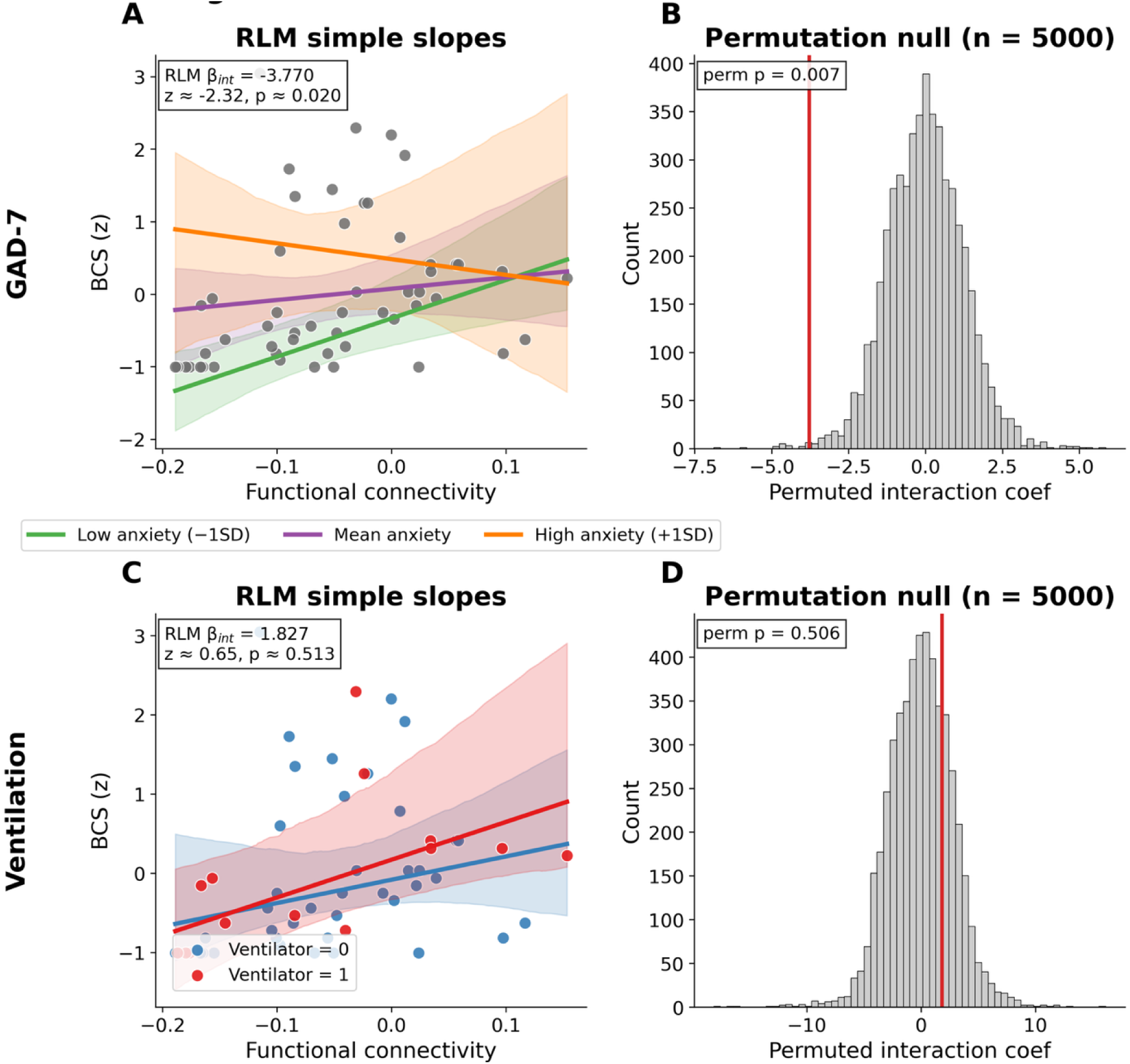
Moderation of the BLA–dACC–BCS association, plotted as in Figure 4. (A, C) RLM simple slopes at representative moderator levels (GAD-7, A; ventilation, C) with 95% case-resampling bootstrap CIs (2,000 resamples) and individual participants. (B, D) Permutation null distributions (N = 5,000) of the connectivity × moderator interaction; the vertical red line marks the observed coefficient.

#### Mechanical ventilation

There was little evidence that mechanical ventilation moderated the BLA–dACC–BCS association (interaction β = 1.83, p = .51; permutation p = .51; Figure 5, bottom row).

### Formal test of differential moderation between edges

The two edges showed different patterns of nominal moderation, which could tempt a double-dissociation interpretation. To test this directly rather than inferring dissociation from one significant and one non-significant test (Nieuwenhuis et al., 2011), we computed post-hoc bootstrap contrasts between the across-edge differences in interaction coefficients (5,000 case-resamples, seed = 42; robust linear regression (RLM, Huber’s T, c = 1.345)). This contrast was prompted by the observed pattern and was not specified in advance. The full-sample interaction coefficients were directionally consistent with a double dissociation: dPAG–PoI1 was more strongly modulated by ventilation (β = −7.24) than by GAD-7 (β = −0.15), whereas BLA–dACC was more strongly modulated by GAD-7 (β = −3.77) than by ventilation (β = 1.83). However, both contrasts included zero: Δ(GAD-7) = −3.62, 95% CI [−9.95, +1.82]; Δ(Ventilation) = −9.07, 95% CI [−21.52, +0.49]. Per the pre-specified decision rule, we therefore do not claim a statistical dissociation between the two edges. The point estimates were sizeable and directionally consistent with separable mechanisms, but the bootstrap distributions were too wide to exclude zero at the present sample size.

### Cohort structure and site-related sensitivity analyses

Recruitment site was partially confounded with clinical subgroup: the Oxford sample comprised 19 previously hospitalised and 9 non-hospitalised participants, while the Cardiff sample comprised 25 non-hospitalised participants exclusively. The three sub-cohorts differed on age (H = 19.49, p < .001), sex distribution (χ^2^ = 17.11, p < .001), days since acute infection (H = 11.02, p = .004), and absolute head motion (H = 6.10, p = .047; controlled in the primary model). Crucially, neither breathlessness outcome differed across sub-cohorts (BCS H = 4.34, p = .114; D-12 H = 3.04, p = .219), indicating comparable breathlessness burden despite differences in clinical phenotype between sites. Connectivity in the dPAG–PoI1 edge did not differ across groups (H = 0.17, p = .920). Connectivity in the BLA–dACC edge did differ across groups (H = 7.60, p = .022), driven primarily by Oxford non-hospitalised versus Cardiff non-hospitalised participants (U = 49.0, Bonferroni-corrected p = .042), which is why this edge was examined under additional covariate specifications. Full cohort characteristics are reported in Appendix 7.

To assess whether the two primary associations were attributable to this cohort structure, we re-fitted each edge model under six alternative covariate specifications, adding (i) site, (ii) three-level cohort group, (iii) the WHO Clinical Progression Scale (WHO severity; Marshall et al., 2020), (iv) days since infection (N = 51), (v) pre-existing respiratory condition (N = 41), and (vi) a reduced covariate set (Table 1). Both associations were preserved across all specifications. In the full sample (N = 53), β for dPAG–PoI1 ranged from −4.38 to −4.74 (all p < .001) and for BLA–dACC from 3.91 to 4.18 (all p ≤ .003). Adjusting for pre-existing respiratory condition in the subsample with available data (N = 41) left both coefficients essentially unchanged relative to the base model refitted in the same subsample (dPAG–PoI1: −3.68 vs −4.00, p = .007; BLA–dACC: 3.58 vs 4.06, p = .039), indicating that the modest attenuation from the full-sample estimates reflects the smaller sample rather than the covariate itself. Neither association was attributable to cohort membership, and hence prior hospitalisation, time since acute infection, pre-existing respiratory disease, motion, or scanner.

**Table 1.**
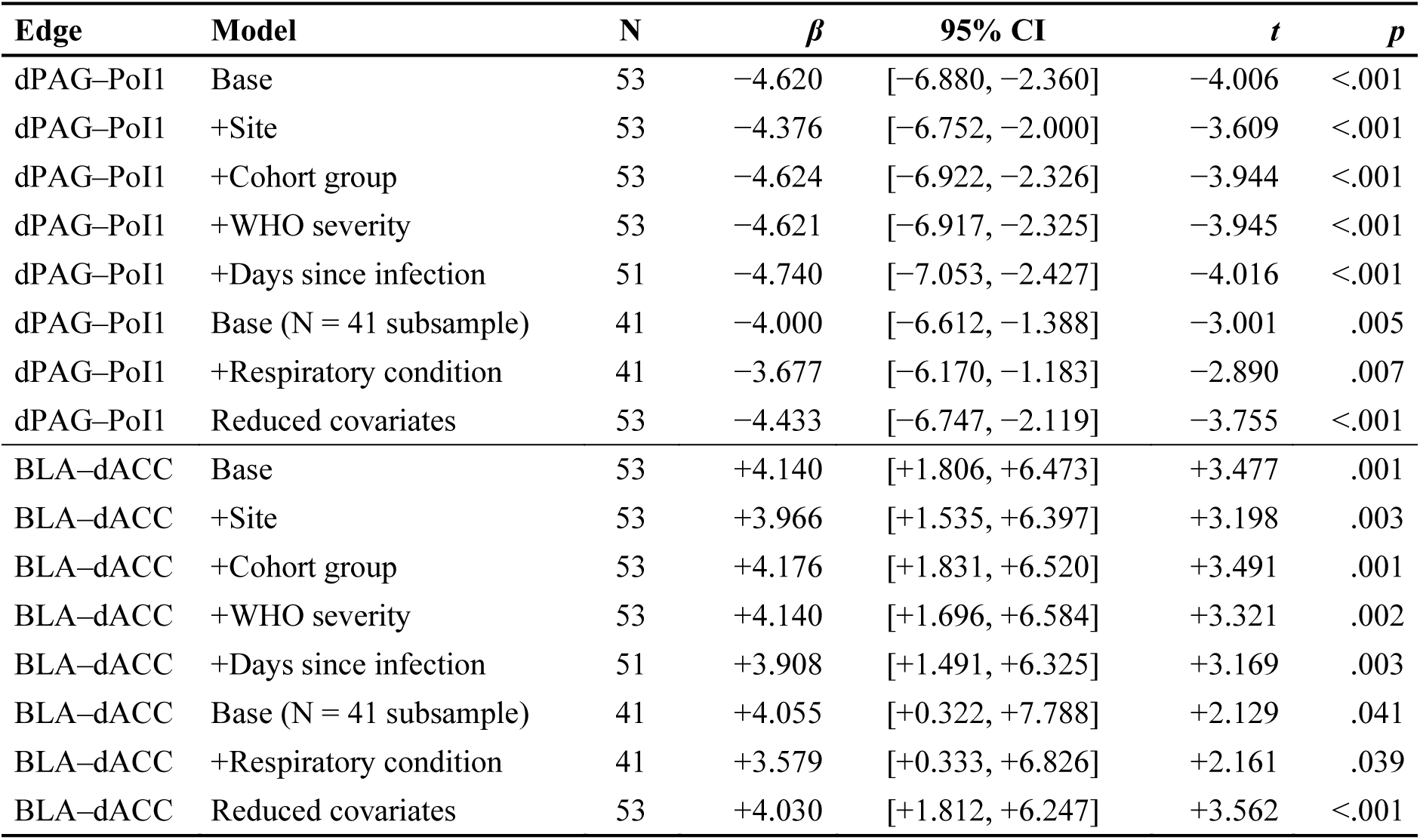
Sensitivity of the two BCS-associated edges across six model specifications (robust linear regression; β [95% CI], p). Models adding days since infection use N = 51; models adding pre-existing respiratory condition use N = 41 (the subsample with available medical history), with the base model refitted in the same subsample for comparison.

## Discussion

In this study, we used 7-tesla resting-state fMRI to examine intrinsic connectivity among 18 regions of interest spanning the interoceptive and visceromotor components of an interoceptive– allostatic framework of respiratory perception and regulation in a post-COVID cohort. Two connections were associated with breathlessness catastrophising (Figure 6). Connectivity between the dorsal periaqueductal grey and posterior insula (dPAG–PoI1) was lower in individuals with greater catastrophising, whereas connectivity between the basolateral amygdala and dorsal anterior cingulate cortex (BLA–dACC) was higher. These two connections showed different patterns of moderation: the dPAG–PoI1 association was stronger in participants who had received mechanical ventilation during acute infection, but showed little evidence of moderation by generalised anxiety, whereas the BLA–dACC association was attenuated at higher levels of generalised anxiety, with little evidence of moderation by mechanical ventilation. Formal contrasts did not, however, establish a statistical dissociation between these moderation patterns. Together, these findings link altered interoceptive signalling and affective threat processing within a single interoceptive– allostatic architecture. Because connectivity was measured at rest rather than during an evoked challenge, these associations index the tonic state in which persistent breathlessness is experienced, rather than the phasic response characterised in experimental paradigms.

**Figure 6.**
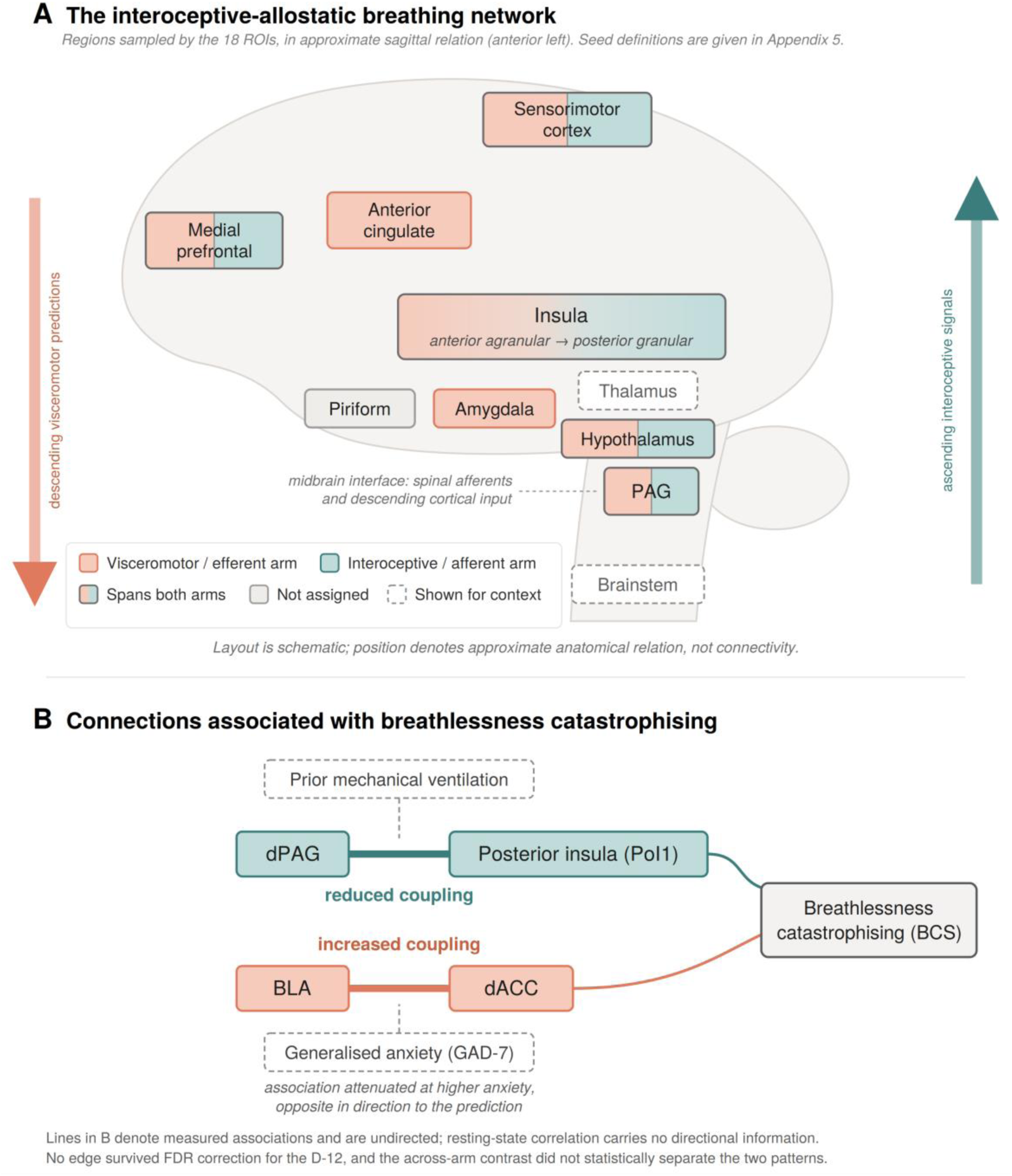
The interoceptive–allostatic breathing network and connections associated with breathlessness catastrophising. (A) Regions sampled by the 18 ROIs, in approximate sagittal relation (anterior left). Colour denotes provisional assignment to the visceromotor (efferent) or interoceptive (afferent) arm following Kleckner et al. (2017), Barrett and Simmons (2015) and Luettich et al. (2023); regions spanning both arms are shown split, and the insula gradient reflects its anterior agranular to posterior granular organisation. The PAG is shown as a midbrain interface rather than a component of either arm. The thalamus and brainstem convey the closed-loop architecture but were not sampled. Flanking arrows indicate proposed direction of information flow; no projections between regions are asserted. Seed definitions, coordinates and lamination are given in Appendix 5. (B) The two edges surviving FDR correction. Dashed leaders indicate the moderator of each association. Lines are undirected. No edge survived correction for the D-12, and the across-arm contrast did not statistically separate the two patterns. The schematic is not lateralised.

### Brainstem–insula hypoconnectivity and the appraisal of breathing

Connectivity between the dorsal periaqueductal grey and the posterior insula (dPAG–PoI1) was negatively associated with breathlessness catastrophising. As the only connection on the interoceptive arm of the framework to survive correction (Figure 6), we treat it as a localised candidate correlate of altered interoceptive signalling rather than evidence for a network-wide effect. The dorsal periaqueductal grey receives spinal sensory afferents and is central to the coordination of respiratory and defensive responses, including perception of dyspnoea (Craven, 2011; Keay and Bandler, 2001; Faull et al., 2019), while the granular posterior insula has been proposed as a primary cortical locus for interoceptive representation and is consistently implicated in breathlessness perception (Kleckner et al., 2017; Herigstad et al., 2011). Within contemporary allostatic–interoceptive models, this pathway may contribute to the integration and weighting of the interoceptive evidence used to infer the state of the body (Barrett and Simmons, 2015; Seth and Friston, 2016). Reduced dPAG–PoI1 coupling may therefore reflect a diminished influence or reliability of ascending respiratory signals, increasing uncertainty about bodily state and limiting the extent to which ongoing sensory evidence can update threat-related expectations about breathing. Benign respiratory sensations may consequently be less readily recognised as signals of safety and more readily interpreted as evidence of persistent respiratory vulnerability.

In the context of post-COVID breathlessness, such a mechanism may bear specifically on how respiratory threat expectations are updated following recovery from acute illness. During acute infection, heightened respiratory vigilance is likely to be adaptive, reflecting genuine physiological threat and a need for increased respiratory preparedness. As respiratory function recovers, interoceptive signals would ordinarily be expected to support the attenuation or extinction of these threat-related expectations. Reduced dPAG–PoI1 coupling may limit the influence of such recovery-related signals, allowing learned beliefs about respiratory vulnerability to persist beyond resolution of the acute threat. On this view, breathlessness catastrophising may reflect a reduced capacity to update or disengage from previously adaptive threat models of breathing.

The stronger association in participants who had required mechanical ventilation during acute infection is consistent with the possibility that this pathway is particularly sensitive to prior respiratory adversity. One candidate mechanism is that prolonged perturbation of respiratory neural activity during critical illness, or the experience of mechanical ventilation itself, induces plasticity within brainstem and spinal respiratory networks that alters the calibration of afferent respiratory signalling (Mahamed et al., 2011; Strey et al., 2013; Albaiceta et al., 2021). Consistent with this, a substantial proportion of invasively ventilated patients report moderate to severe breathlessness during ventilation (Demoule et al., 2024), and brainstem alterations have been reported following severe COVID-19 (Rua et al., 2024).

Affective contributions are nonetheless likely. Intensive care admission and mechanical ventilation carry a substantial psychological burden, and post-traumatic symptoms are common among survivors of critical respiratory illness (Davydow et al., 2008; Parker et al., 2015). The GAD-7 used here indexes generalised anxiety over the two weeks preceding the scan and does not capture trauma-related distress or breathing-specific fear. The lack of evidence for GAD-7 moderation therefore constrains only the contribution of contemporaneous generalised anxiety and does not exclude a broader affective legacy of critical illness. Residual pulmonary insult, trauma-related distress, and altered interoceptive processing are likely to co-occur in previously ventilated participants, and the present design cannot separate them. We therefore regard the dPAG–PoI1 pathway as a candidate lower-level component of interoceptive regulation, through which acute respiratory insults may exert lasting influence on the perception and appraisal of breathing, alongside affective and physiological consequences that our measures were not designed to capture.

### Limbic–cingulate hyperconnectivity: threat appraisal coupled to visceromotor control

Connectivity between the basolateral amygdala and the dorsal anterior cingulate cortex (BLA– dACC) was positively associated with breathlessness catastrophising. As the only connection on the visceromotor arm of the framework to survive correction (Figure 6), we interpret it as a localised candidate correlate of altered threat-related processing and visceromotor control rather than evidence for a network-wide abnormality. The BLA is thought to encode the affective significance of sensory and interoceptive cues, supporting the acquisition and expression of learned threat associations (LeDoux, 2000; Herry and Johansen, 2014; Janak and Tye, 2015), while the dACC has been implicated in threat appraisal, the anticipation of aversive outcomes, and the regulation of behavioural and autonomic responses to biologically salient challenges (Critchley et al., 2004; Shackman et al., 2011; Barrett and Simmons, 2015). Within contemporary allostatic– interoceptive models, the agranular dACC is regarded as a key visceromotor region that generates predictions about bodily needs and coordinates the regulatory responses supporting physiological stability (Barrett and Simmons, 2015). Stronger coupling between these regions may therefore indicate that threat-related interpretations of respiratory sensations are more readily translated into anticipatory visceromotor responses.

Rather than reflecting a purely perceptual bias, this pattern is consistent with a closed-loop account of interoceptive processing in which predictions about bodily threat are continuously linked to action-oriented regulatory responses. Such a view accords with interoceptive–allostatic accounts of brain function and with theories of constructed emotion, which hold that affective experience emerges through ongoing interactions between predictions about bodily state and the regulatory processes that maintain physiological stability (Barrett and Simmons, 2015; Barrett, 2017). It also echoes William James’s notion of the “subtler emotions”, the diffuse and often unconscious bodily feelings that shape perception and behaviour without requiring explicit emotional awareness (James, 1948). On this account, BLA–dACC coupling may index a persistent allostatic bias towards respiratory threat, characterised by heightened physiological preparedness and increased sensitivity to ambiguous interoceptive input. This construct sits at the intersection of several existing accounts: allostatic load and anticipatory regulation (McEwen, 2003; Sterling, 2012), anxiety sensitivity and fear of bodily sensations (Reiss, 1985; Taylor, 2014; Paulus and Stein, 2010), and interoceptive predictive-processing models in which strong threat priors dominate ascending sensory evidence (Seth et al., 2012; Allen and Tsakiris, 2018). Such a bias could represent either a pre-existing vulnerability or a learned adaptation acquired during acute illness, whereby repeated experiences of respiratory distress strengthen associations between bodily sensations and threat.

This association was moderated by generalised anxiety scores, with little evidence of moderation by mechanical ventilation, the reverse of the moderation profile observed for dPAG–PoI1. Whereas the dPAG–PoI1 association appeared more closely tied to respiratory adversity, the moderation of BLA–dACC connectivity is more dependent on higher-order affective and cognitive processes, in line with longstanding accounts that emphasise the contribution of anxiety, threat appraisal, and emotional response to symptom amplification (Bailey, 2004; Hayen et al., 2013; Faull et al., 2019). The direction of the effect, however, was not what we had predicted: the association between BLA–dACC connectivity and catastrophising was strongest at lower levels of anxiety and attenuated at higher levels. This is difficult to reconcile with a simple anxiety-driven account. Were BLA–dACC connectivity merely the neural expression of generalised anxiety, its association with catastrophising should have been strongest in the most anxious individuals; the opposite was observed.

One possibility is that BLA–dACC connectivity reflects a threat-processing mechanism relatively specific to breathlessness rather than a nonspecific marker of negative affect. On this view, catastrophising arises from learned expectations or threat beliefs about respiratory sensations that are partially separable from broader anxiety levels, consistent with evidence that dyspnoea-related fear and catastrophising relate more specifically to respiratory symptom perception than to negative affectivity in general (Willgoss et al., 2013; Janssens et al., 2011). When generalised anxiety is low, variation in catastrophising may be governed more directly by this respiratory-specific threat processing. At higher levels of anxiety, catastrophising may be shaped by a broader range of affective and cognitive influences, reducing the relative contribution of BLA–dACC coupling. Generalised anxiety therefore seems unlikely to provide a complete explanation for this association and may instead modulate the extent to which breathlessness-specific threat representations are expressed. BLA–dACC coupling may thus reflect a breathlessness-related threat–visceromotor process that is modulated, rather than simply explained, by generalised anxiety.

### An integrative account of breathing as an interoceptive–allostatic perception–action loop

Taken together, the reduced dPAG–PoI1 and increased BLA–dACC connectivity offer a candidate account of symptom–biomarker discordance in the perception of breathing. Rather than a simple mismatch between pulmonary physiology and subjective experience, breathlessness catastrophising may reflect variability in how interoceptive signals are integrated with threat-related expectations within a closed-loop system that governs perception and physiological regulation jointly. Within such a loop, respiratory physiology, anxiety and beliefs about breathing are not necessarily independent contributors: threat-related expectations may shape respiratory and autonomic regulation, whose interoceptive consequences can in turn reinforce the experience of breathlessness. Priors shaped during acute illness could therefore influence respiratory output as well as respiratory experience. Autonomic dysregulation is well recognised following viral infection, most clearly as inappropriate sinus tachycardia (Dani et al., 2021; Eastin et al., 2025; Keller et al., 2026). A respiratory counterpart might involve altered breathing pattern, such as elevated respiratory rate, greater breath-to-breath variability or a shift towards upper-thoracic breathing, rather than an altered interpretation of an otherwise unchanged pattern. On this account, dysfunctional breathing and persistent breathlessness need not be competing explanations but may represent complementary manifestations of a dysregulated perception–action loop: one expressed in ventilatory behaviour and the other in subjective experience.

The different moderation patterns provide tentative support for partially separable influences within this system: the dPAG–PoI1 association was more strongly related to acute respiratory severity, whereas the BLA–dACC association varied with generalised anxiety. However, formal across-edge contrasts did not statistically distinguish these moderation effects. We therefore regard them as suggestive of potentially separable influences rather than evidence for a double dissociation, and the perception–action account presented here as a framework for interpreting the observed findings rather than a demonstrated causal architecture.

### Catastrophising versus global symptom burden

The two FDR-corrected associations were observed for the BCS (Solomon et al., 2015) rather than the D-12 (Yorke et al., 2010). The BCS indexes threat-related appraisal of respiratory sensations, whereas the D-12 captures broader sensory and affective symptom burden. We had anticipated that these measures would preferentially map onto different arms of the network, with catastrophising associated most closely with visceromotor and limbic coupling and D-12 with interoceptive coupling. This measure-level separation was not observed. Both surviving edges, one from each arm of the loop, were associated with catastrophising, whereas no D-12 edge survived correction.

Rather than distinguishing the two pathways, the measures appeared to differ in their overall sensitivity to the connectivity patterns examined here. The uncorrected D-12 coefficients closely resembled those for catastrophising, with positive associations concentrated mainly among visceromotor connections and negative associations among interoceptive connections. Although these associations did not survive correction, this similarity raises the possibility that the difference between BCS and D-12 reflects phenotypic specificity or statistical sensitivity rather than wholly distinct neural mechanisms.

### Strengths and limitations

This study combined ultra-high-field 7 T resting-state fMRI with an extended acquisition enabling more sensitive measurement of connectivity involving small brainstem structures and fine-grained insular subdivisions that are difficult to resolve at conventional field strengths. Inference was constrained to an a priori interoceptive–allostatic network of breathing, reducing analytic flexibility relative to unconstrained whole-brain searches. Robustness was assessed using FDR-controlled robust regression, leave-one-out and bootstrap diagnostics, and sensitivity analyses across alternative cohort and covariate specifications.

Several limitations warrant consideration. The modest sample size restricts stable multivariable modelling of the full range of potential clinical confounds and, in particular, the precision of the exploratory moderation analysis. Site, recruitment pathway, hospitalisation, and mechanical ventilation were structurally confounded. All 14 ventilated participants had been hospitalised and scanned at Oxford, whereas no ventilated participants were recruited at Cardiff. Although the two primary edge associations were preserved after adjustment for site, cohort group and acute severity, these sensitivity analyses concerned the main associations rather than the moderation effects. Mechanical ventilation occurred alongside other features of severe critical illness, including hypoxaemia, sedation, prolonged immobility, and intensive-care exposure, which cannot be disentangled in the present sample. The generalised anxiety moderator (GAD-7) was right-skewed, with relatively few participants at higher anxiety levels, making estimates in this range less precise. The leave-one-out and secondary moderation analyses were conducted on already-selected edges and do not constitute independent validation.

Pulmonary physiology was not systematically characterised. Spirometry was available in 33 participants and was within normal limits in those measured, consistent with the broader post-COVID literature summarised in the Introduction. Spirometry is, however, relatively insensitive to subtle abnormalities, and diffusion limitation, small-airways dysfunction, dysfunctional breathing, deconditioning and autonomic dysregulation cannot be excluded. The present findings should therefore be read as characterising breathlessness that is disproportionate to any demonstrable pulmonary impairment, rather than as excluding peripheral contributions altogether.

A related concern is whether the observed connectivity patterns are specific to post-COVID breathlessness or reflect chronic respiratory disease more broadly. Information on pre-existing respiratory and cardiac conditions was available for 77% of participants. Respiratory history was associated with breathlessness severity but not with either connectivity edge, and adjusting for it in a sensitivity model preserved both primary associations. A more conservative analysis restricted to participants without respiratory history retained the direction of both effects but was underpowered at N = 31. More complete pulmonary and clinical phenotyping in future cohorts would be needed to substantiate this interpretation.

The BCS, D-12 and GAD-7 are self-report measures and therefore depend on participants’ subjective appraisal and recall. Scores may vary with contemporaneous affect and day-to-day symptom fluctuations, while between-person differences in response style may introduce additional measurement variability. Repeated assessments and complementary behavioural or physiological measures would provide a more direct characterisation of the processes inferred here.

The cross-sectional design precludes inference about whether altered connectivity predisposes to persistent breathlessness, arises following acute illness or persistent symptoms, or reflects a combination of these processes. Resting-state functional connectivity is also an indirect and undirected measure of neural coupling, and analyses of small brain structures are particularly susceptible to physiological noise and susceptibility artefacts despite extensive denoising. ROI localisation based on standard-space templates may additionally be affected by individual anatomical variability, particularly in small brainstem nuclei.

Both sites used the same model of Siemens Magnetom 7 T scanner and followed the UK7T Network’s harmonised anatomical and functional imaging protocols (Clarke et al., 2020), designed to minimise inter-scanner differences. The UK7T Travelling Heads study nevertheless reported lower inter-than intra-site reproducibility of 7 T fMRI, although the magnitude of these differences were small and comparable with published single-site estimates (Driver et al., 2021). In the present data, adjustment for site attenuated the two primary edge coefficients by <6% (Table 1), arguing against a simple between-site offset as an explanation for either association. Harmonisation cannot, however, remove differences in clinical sampling between centres, and site remained partly confounded with hospitalisation and ventilation status.

### Future directions

Future work should replicate these findings in larger, clinically balanced cohorts and combine longitudinal neuroimaging with respiratory challenges and concurrent ventilatory and autonomic measurements. Such designs could test whether altered interoceptive processing and visceromotor regulation predict symptom persistence or recovery, and whether similar mechanisms generalise to other conditions characterised by disproportionate breathlessness. Computational models of interoceptive inference could further help quantify individual differences in the weighting of respiratory evidence and threat-related priors, providing a more direct test of the mechanisms proposed here.

### Conclusion

In this study we identified two resting-state connections associated with breathlessness catastrophising in post-COVID patients: reduced functional connectivity between dorsal periaqueductal grey and posterior insula and increased functional connectivity between basolateral amygdala and dorsal anterior cingulate. The two associations showed different patterns of moderation by acute respiratory severity and generalised anxiety, although formal comparisons did not establish a statistical dissociation, and no connectivity association with D-12 survived correction.

The two edges occupy complementary interoceptive and visceromotor components of the proposed respiratory perception–action loop. We propose that persistent post-COVID breathlessness may involve altered coordination between interoceptive evidence and threat-related respiratory regulation, rather than perceptual amplification alone. This is a first step towards an account of persistent breathlessness that locates the disturbance in how breathing is regulated as much as in how it is perceived, and one that is neither purely physiological nor dismissively psychological.

## Materials and methods

### Participants

A total of 53 patients (59% female; mean age 50 ± 14 years; mean BMI 28 ± 5) were scanned a median of 257 days (IQR 179–371; range 125–806; mean 323 ± 196) after COVID-19 infection. All participants underwent 7 tesla MRI scanning at one of two sites: the Oxford Centre for Integrative Neuroimaging (OxCIN; formerly the Wellcome Centre for Integrative Neuroimaging [WIN]; Oxford, UK) and the Cardiff University Brain Research Imaging Centre (CUBRIC; Cardiff, UK).

At OxCIN, 28 patients were scanned, comprising 19 individuals who had been hospitalised and subsequently discharged from the John Radcliffe Hospital (Oxford, UK) and 9 non-hospitalised patients referred by the Oxford Post-COVID Service at the Churchill Hospital (Oxford, UK). At CUBRIC, 25 non-hospitalised patients referred by the Cardiff and Vale University Health Board (Cardiff, UK) were scanned.

Recruitment began with telephone screening to assess eligibility for 7 T MRI, with primary emphasis on safety compliance for referred patients. Following informed consent, participants completed pre-scan questionnaires electronically via a secure platform (JISC). An exception was made for the initial Oxford cohort (N = 12), who completed paper-based questionnaires owing to the rapid initiation of scanning during the evolving circumstances of the pandemic and the concurrent development of the online survey system. All questionnaires, including a cognitive task not analysed in the present study, were completed within five days of MRI acquisition.

The study received approval from the North West – Preston Research Ethics Committee. All participants provided informed consent either for the sub-study CMORE-NEURO of the CMORE project, titled Assessing the Effects of Coronavirus Disease (COVID-19) on Multiple Organ Systems and Impact on Quality of Life, Functional Capacity, and Mental Health (REC reference: 20/NW/0235; IRAS project ID: 282608), or for the BBB-COV study, titled Brain and Brainstem Basis of Persistent Symptoms in COVID-19 (REC reference: 21/NW/0090; IRAS project ID: 297314). The study was not pre-registered.

### Behavioural measures

Post-COVID breathlessness was assessed using two complementary self-report measures capturing distinct facets of the breathlessness experience. The Breathlessness Catastrophising Scale (BCS; Solomon et al., 2015), adapted from the Pain Catastrophising Scale, indexes threat-related appraisal of respiratory sensations, that is, the tendency to interpret breathing difficulty in catastrophic or highly negative terms. The Dyspnoea-12 (D-12; Yorke et al., 2010) captures a broader, multidimensional profile of symptom burden, spanning both the sensory and affective dimensions of breathlessness. Together, these instruments distinguish maladaptive appraisal of breathing from overall symptom severity. Generalised anxiety scores were measured using the Generalised Anxiety Disorder 7-item scale (GAD-7; Spitzer et al., 2006; Kroenke et al., 2010). Further details of all behavioural measures are provided in Appendix 3.

Demographic information, medical history, and clinical data during the acute infection were obtained from hospital records or directly from participants at the time of the study (within five days before the scan). These data included infection date (for computation of days from infection to scan), acute-stage symptoms, markers of inflammation (peak C-reactive protein and D-dimer), and spirometry indices (FEV₁, FVC, FEV₁/FVC). Severity of acute COVID-19 infection was primarily defined by the requirement for mechanical ventilation in intensive care, which by definition indicates respiratory failure. A subset of participants had also been assessed with the WHO Clinical Progression Scale (Marshall et al., 2020); however, data for this scale were incomplete and were not suitable for primary analyses.

### MRI scanning sequences

Resting-state functional data were acquired using the CMRR multiband EPI sequence (Moeller et al., 2010) in a single 10-minute run (400 volumes), with a TR of 1.5 s, TE of 25 ms, isotropic resolution of 1.5 mm, 96 slices and a multiband factor of 4. The EPI volume was positioned to cover the whole brain. A single-band whole-brain high-contrast EPI scan was acquired to aid registration of the functional data, and B0 field maps were acquired for distortion correction. For anatomical localisation and registration, a T1-weighted MP2RAGE was acquired with 0.7 mm isotropic voxels, TE of 2.64 ms, TR of 3500 ms, bandwidth of 300 Hz/pixel, inversion times of 725 and 3150 ms, nominal flip angles of 5° and 2°, and a matrix size of 224 × 224 × 224. Image processing used routines from the FMRIB Software Library (FSL) v6.0.7.6 (Jenkinson et al., 2012), Advanced Normalization Tools (ANTs) v2.2.0 (Avants et al., 2011), SPM12 v7219 (Penny et al., 2011), MATLAB R2022b and Python 3.11.6. Both sites are members of the UK7T Network, and acquisition followed the Network’s harmonised anatomical and functional protocols (Clarke et al., 2020).

### Brain imaging data processing

#### Anatomical data preprocessing

MP2RAGE images from both inversion times were combined via Phase-Sensitive Inversion Recovery (PSIR; Clarke et al., 2020). A liberal brain-extraction mask was applied to facilitate registration, followed by bias-field correction and segmentation into grey matter, white matter, and cerebrospinal fluid (CSF).

#### Resting-state EPI preprocessing and registration

Resting-state fMRI data underwent brain extraction, motion correction, high-pass temporal filtering (cut-off 100 s; Smith et al., 2013), and distortion correction using FUGUE and boundary-based registration. Data were linearly registered (6 degrees of freedom) to the anatomical image via FLIRT and nonlinearly transformed to MNI152 standard space (1-mm resolution) using FNIRT. Spatial smoothing was omitted during preprocessing to preserve subject-specific independent components, though a minimal 1.5-mm kernel was applied during final FEAT analysis.

#### Physiological and ICA-based denoising

Physiological noise correction and structured artefact removal were implemented using a combined ICA–PNM pipeline. ICA+FIX was applied via MELODIC to decompose the 4D EPI data into spatial and temporal components, with a manually labelled training dataset (N = 20) balanced across hospitalisation status and acquisition site informing classifier training (Griffanti et al., 2014). Simultaneously, physiological data (cardiac and respiratory signals) acquired alongside EPI triggers, using BIOPAC (Oxford) and an ADI setup (Cardiff), were processed via FSL’s PNM tool (Glover et al., 2000). Phase information was modelled using a low-order Fourier expansion, generating regressors for respiratory (N = 4), cardiac (N = 3), and interaction (N = 1) components (Brooks et al., 2008; Harvey et al., 2008; Faull and Pattinson, 2017). PNM stage-1 outputs were visually inspected before subsequent stages. The resulting denoising model incorporated identified ICA noise components, subject-specific white-matter and CSF time series, and the PNM regressors within a unified GLM in FEAT. This integrated approach mitigates variance reintroduction from overlapping noise sources, yielding substantially improved temporal signal-to-noise ratio (Appendix 4). The output is referred to as the ‘cleaned data’ throughout.

#### ROI identification

We identified 18 ROIs believed to be central to the interoceptive and allostatic network associated with respiration (Kleckner et al., 2017; Hassanpour et al., 2018; Luettich et al., 2023). Regions within ACC and insula were defined using the Harvard–Oxford cortical and subcortical atlases and the Glasser atlas (Glasser et al., 2016), with refinement based on 4-mm spheres centred on MNI coordinates from Kleckner et al. (2017). When precise seeds overlapped with broader atlas-derived masks, the 4-mm sphere seeds were retained for specificity. Additional regions not covered by Kleckner et al. (2017) but falling within the broader ACC or insular domains were included to ensure full anatomical coverage. Selected ROIs included the subgenual, pregenual, and dorsal ACC (each 4 mm), and key insular subregions: medial and lateral ventral anterior insula, dorsal medial insula, dorsal posterior insula (all 4 mm), and posterior insular areas PoI1 and PoI2 (from the Glasser atlas). The anterior insula comprised AVI and AAIC (agranular); the middle insula was represented by MI (dysgranular); and the posterior insula included PoI1 and PoI2. The granular Ig region, overlapping conceptually with the dpIns seed, was excluded to avoid redundancy. The dorsal PAG (dPAG), ventrolateral PAG (vlPAG), and basolateral amygdala (BLA) ROIs were derived from masks used in prior breathlessness-related studies (Faull et al., 2016; Faull and Pattinson, 2017; Qiu, 2025).

The 2-mm ‘lPAG’ seed reported in earlier breathlessness work (Faull et al., 2016; Faull and Pattinson, 2017) is encompassed within the current dPAG mask. The hypothalamus ROI was individually segmented using the FreeSurfer Hypothalamic Subunits tool (v7.2; Billot et al., 2020). Piriform cortex masks were adapted from Zhou et al. (2019). The S1, M1, vmPFC, and dmPFC ROIs were obtained using the Harvard–Oxford cortical atlas. A summary of the ROIs, with lamination characteristics (Kleckner et al., 2017) and putative function, is provided in Appendix 5.

#### FC extraction

ROI masks were transformed into each subject’s functional EPI space, followed by quality-control procedures to verify mask validity (volume, anatomical location, and signal-to-noise ratio; Appendix 6). Mean time courses were extracted for each ROI in subject-specific EPI space, generating a matrix of dimensions 18 × 400 × 53 (ROIs × time points × subjects). Functional connectivity between all ROI pairs (edges) was computed as Pearson correlation coefficients, producing a final connectivity matrix of size 53 × 153 (subjects × unique edges, derived from pairwise combinations of 18 nodes).

### Associating FC measures with behavioural outcomes

To evaluate associations between functional connectivity and post-COVID breathlessness, univariate robust linear regression models (RLM, Huber’s T, c = 1.345) were implemented in Python 3.11.6 using statsmodels. Breathlessness was indexed using the Dyspnoea-12 (Yorke et al., 2010) and the Breathlessness Catastrophising Scale (Solomon et al., 2015). The BCS and D-12 outcomes were standardised (z-scored) prior to analysis. Robust regression reduces sensitivity to outliers and accommodates potential non-normality and heteroscedasticity in residuals. D-12 and BCS were modelled separately as dependent variables, with each of the 153 edges entered in turn as the predictor of interest, adjusted for age, sex, mean absolute and relative framewise displacement, BMI, brain volume, and intracranial volume. Benjamini–Hochberg FDR correction (Benjamini and Hochberg, 1995) was applied across the 153 edges within each outcome.

#### Covariate selection rationale

The primary model was kept parsimonious because the sample size (N = 53) was modest relative to the number of plausible demographic, clinical, and imaging covariates. Covariates were specified a priori for their established influence on resting-state functional connectivity estimates or breathlessness-related reporting (age, sex, BMI, absolute and relative mean framewise displacement, cube-root-transformed total intracranial volume, and cube-root-transformed total brain volume) rather than on the basis of univariate associations with the outcome. We explicitly did not interpret non-significant univariate associations as evidence that a candidate variable was unimportant, since a covariate can confound an edge–outcome relationship even when its marginal association with the outcome is null.

Clinical variables that were incomplete or strongly collinear with site/cohort structure were not entered in the primary model. Site, cohort group (which nests hospitalisation status), acute severity on the WHO Clinical Progression Scale, days since infection and pre-existing respiratory condition were instead examined in the sensitivity analyses (Table 1). Ventilation status was examined as a moderator rather than as a covariate, and pulmonary-function measures were not entered in any model owing to missingness (Appendix 8). Blood pressure was excluded owing to substantial missing data and inconsistent measurement timing. Full rationale for all 16 candidate covariates, with their univariate associations and missingness, is provided in Appendix 8.

One inclusion decision warrants explicit comment. Generalised anxiety scores (GAD-7) showed the largest univariate association with BCS of any non-imaging variable (rₛ = .364, p = .007), yet were deliberately excluded from the base model because GAD-7 was uncorrelated with either FC edge (rₛ = .033 for dPAG–PoI1; rₛ = .038 for BLA–dACC). This is the pattern expected if anxiety operates downstream of the connectivity–breathlessness relationship rather than upstream as a confounder. Conditioning on a downstream mediator would attenuate a genuine brain–behaviour association (overadjustment). Because cross-sectional data cannot definitively distinguish a confounder from a mediator, GAD-7 was reserved for the moderation analysis rather than entered as a base covariate.

#### Moderation and dissociation analyses

Moderation analyses were secondary and exploratory and were conducted only for the two edges that survived FDR correction in the BCS analysis. For each edge, interaction terms with GAD-7 and with mechanical ventilation status were tested using robust linear regression (RLM, Huber’s T, c = 1.345) and the primary covariate set. Non-parametric permutation tests (N = 5,000) provided empirical p-values. To test whether moderation effects differed between the two edges rather than relying on parallel patterns of significance (Nieuwenhuis et al., 2011), we computed bootstrap contrasts between the across-edge differences in interaction coefficients (5,000 case-resamples, seed = 42). We pre-specified that a 95% percentile confidence interval excluding zero would be interpreted as evidence that the moderation effect differed between edges, while a confidence interval including zero would lead us to drop any claim of statistical dissociation.

#### Internal-stability diagnostics

To characterise the sensitivity of the two FDR-surviving edge associations to individual observations, we computed leave-one-out (LOO) coefficient estimates, per-observation influence diagnostics (DFBETA, DFBETAS, Cook’s distance, leverage), and bias-corrected accelerated (BCa) bootstrap 95% confidence intervals from 5,000 case-resampling iterations (seed = 42), with the LOO estimates serving as the jackknife acceleration input. Cook’s distance was computed using the OLS hat matrix and OLS residual variance rather than the RLM robust scale, because the Huber robust scale is systematically smaller than the OLS sigma and would inflate Cook’s distance (by a factor of approximately seven for these models); this choice is documented in the analysis logfile. These analyses are reported as internal-stability checks on the two pre-selected associations and are not intended as independent validation.

## Acknowledgements

We thank the participants who took part in this study, and the clinical teams at the John Radcliffe Hospital, the Oxford Post-COVID Service at the Churchill Hospital, and the Cardiff and Vale University Health Board for their support with recruitment.

This work was supported by the NIHR Oxford Biomedical Research Centre (BRC), based at Oxford University Hospitals NHS Foundation Trust and the University of Oxford, and by the NIHR Oxford Health Biomedical Research Centre (NIHR203316). Additional support was provided by the Oxford University Medical Sciences Division COVID-19 Rapid Response Fund. The Cardiff arm of the study was supported by the Welsh Government through the Wales COVID-19 Evidence Centre, Primary Evaluation Activities Programme (reference 522128). WTC is funded by Wellcome (225924/Z/22/Z). BR is funded by a Wellcome Career Development Award (302210/Z/23/Z) and acknowledges support from the NIHR Oxford BRC and the British Heart Foundation Oxford Centre of Research Excellence. SC and the Oxford Centre for Integrative Neuroimaging were supported by core funding from the Wellcome Trust (203139/Z/16/Z and 203139/A/16/Z). The views expressed are those of the authors and not necessarily those of the NIHR or the Department of Health and Social Care.

This research was funded in whole, or in part, by the Wellcome Trust (225924/Z/22/Z, 302210/Z/23/Z, 203139/Z/16/Z and 203139/A/16/Z). For the purpose of open access, the authors have applied a CC BY public copyright licence to any Author Accepted Manuscript version arising from this submission.

