## Appendix for "Breathlessness catastrophising after COVID-19 involves both interoceptive and visceromotor connectivity"

**Appendix 1 — Spirometry Data**

Spirometry was available for a subset of participants (N = 33), predominantly hospitalised individuals. Group-level values were within normal limits (FEV1 %pred: 95 ± 18; FVC %pred: 104 ± 18; FEV1/FVC: 0.79 ± 0.06), consistent with the broader long-COVID literature (Stewart et al., 2023; Evans et al., 2022). One exception was a 76-year-old male with FEV1/FVC = 64.4, which remained within acceptable limits. Owing to the extent of missing data, spirometry was not included as a covariate in the regression models; all participants were considered to have relatively normal lung function.

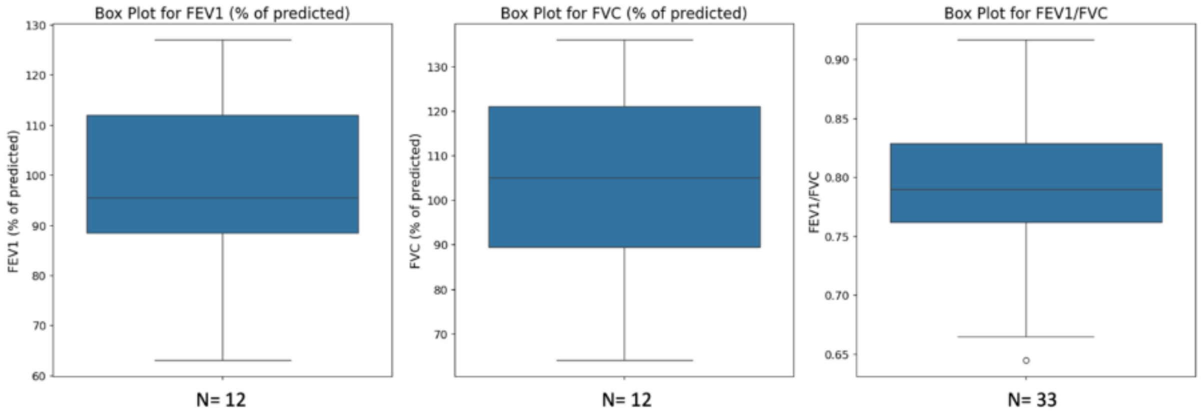


**Appendix 1—figure 1.** Spirometry data for participants who completed pulmonary function testing (N = 33). Boxplots show FEV1 (%pred), FVC (%pred), and FEV1/FVC. Dashed lines indicate lower limits of normal (FEV1/FVC < 0.70; %pred < 80).

### Appendix 2—Demographic Information of Participants

**Appendix 2—table 1.** Demographic information of participants.

| Demographic variable | Value |
|---|---|
| Participants | 53 patients |
| Hospitalisation (count, %) | 19, 36% |
| Age at consent, years (mean, SD, range) | 50 ± 14 (20–79) |
| Gender (female count, %) | 31, 59% |
| BMI (mean, SD, range) | 28 ± 5 (18–43) |
| WHO Clinical Progression Scale (mean, SD, range) | 2.4 ± 2.0 (0–7) |
| FEV1 %pred (mean, SD, range) | 95 ± 18 (61–127) |
| FVC %pred (mean, SD, range) | 104 ± 18 (61–139) |
| FEV1/FVC (mean, SD, range) | 79 ± 6 (64–92) |
| Days from infection to scan (mean, SD, range) | 323 ± 196 (125–806) |

### Appendix 3 — Behavioural Measures

Four validated self-report questionnaires were administered to capture breathlessness severity, cognitive–emotional responses, generalised anxiety scores, and acute infection severity.

#### *Dyspnoea-12 (D-12)*

The D-12 (Yorke et al., 2010) is a 12-item questionnaire measuring the physical and emotional dimensions of breathlessness. Each item is rated on a four-point Likert scale (none to severe); total scores range from 0 to 36, with higher scores indicating greater dyspnoea severity. The D-12 has been validated across multiple respiratory conditions (Yorke et al., 2011; Yorke and Armstrong, 2014; Williams et al., 2017; Simsic et al., 2018).

#### *Breathlessness Catastrophising Scale (BCS)*

The BCS (Solomon et al., 2015) is a 13-item instrument assessing cognitive and emotional responses to breathlessness. Items are rated on a five-point Likert scale (0 = not at all, 4 = all the time), yielding total scores from 0 to 52, with higher scores indicating greater catastrophising. The BCS was adapted from the Pain Catastrophising Scale (Sullivan et al., 1995) and validated in COPD populations (Solomon et al., 2015), with links to breathlessness anticipation demonstrated in fMRI research (Stoeckel et al., 2018).

#### *Generalised Anxiety Disorder 7-item Scale (GAD-7)*

The GAD-7 (Spitzer et al., 2006; Kroenke et al., 2010) is a 7-item self-report measure of anxiety scores frequency over the preceding two weeks. Items are rated 0–3 (not at all to nearly every day), giving a total score of 0–21. Threshold scores of 5, 10, and 15 correspond to mild, moderate, and severe anxiety; a score $\geq 10$ indicates clinically significant anxiety (sensitivity 89%, specificity 82%; Spitzer et al., 2006).

#### *WHO Clinical Progression Scale*

The WHO Clinical Progression Scale (Marshall et al., 2020) is an ordinal measure of acute COVID-19 severity, ranging from 0 (no infection) to 8 (death). It was used here to index acute infection severity at the time of hospitalisation.

**Appendix 4 — Temporal Signal-to-Noise Ratio Improvement After Denoising**

Following combined FIX and PNM denoising, mean tSNR across all ROI masks exceeded 40 in all cases, indicating acceptable signal quality for functional connectivity analysis. Percentage increases in tSNR relative to raw EPI data ranged from 354–607% across ROIs, confirming that the integrated denoising pipeline substantially reduced noise without reintroducing variance from overlapping sources.

**Appendix 4—Table 1.** Percentage increase in tSNR for each ROI mask following FIX and PNM denoising, relative to raw EPI functional data.

| ROI mask | tSNR increase (%) | ROI mask | tSNR increase (%) |
|---|---|---|---|
| vlPAG | 364.59 | lvaIns | 510.14 |
| dPAG | 357.94 | dmIns | 427.57 |
| piri | 484.22 | PoI1 | 399.76 |
| Hyp | 481.74 | PoI2 | 427.34 |
| amyg | 396.71 | dpIns | 354.13 |
| sgACC | 424.97 | dmPFC | 532.65 |
| pACC | 461.86 | vmPFC | 606.59 |
| dACC | 503.30 | S1 | 559.47 |
| mvaIns | 386.28 | M1 | 508.85 |

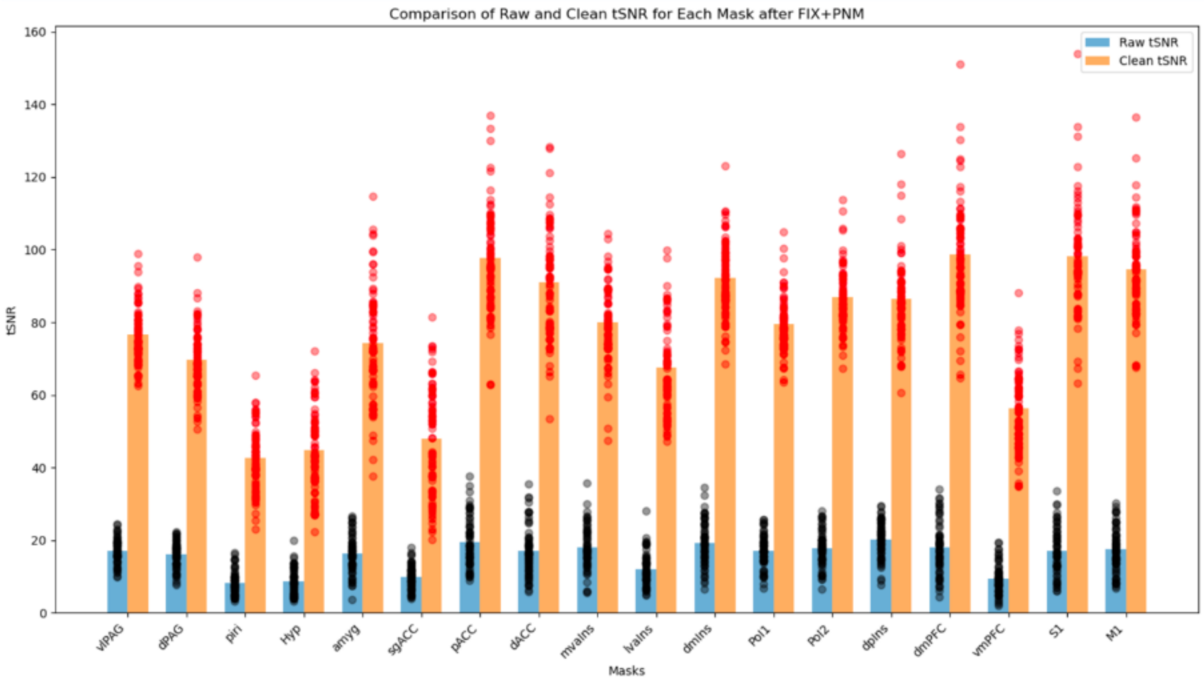


**Appendix 4—Figure 1.** Improvement in tSNR following FIX and PNM denoising across all ROI masks. All ROIs exceeded the minimum acceptable threshold of 40.

### Appendix 5 — Region of Interest Summary

Eighteen ROIs were selected to cover the interoceptive–allostatic breathing network. Subcortical regions comprised the basolateral amygdala (BLA; inherited mask, MNI 22, −3, −19), piriform cortex (4-mm sphere, MNI 16, −2, −14), dorsal and ventrolateral periaqueductal grey (dPAG and vlPAG; inherited masks), and hypothalamus (automated T1w segmentation). ACC regions included the subgenual (sgACC; MNI 2, 14, −6), pregenual (pACC; MNI 13, 44, 0), and dorsal ACC (dACC/aMCC; MNI 9, 22, 33), all 4-mm spheres. Insula regions comprised the dorsal posterior (dpIns; MNI 36, −32, 16), dorsal medial (dmIns; MNI 41, 2, 3), and medial and lateral ventral anterior insula (mvaIns, MNI 30, 16, −14; lvaIns, MNI 44, 6, −15), all 4-mm spheres, plus posterior insula subdivisions PoI1 and PoI2 from the Glasser atlas (Glasser et al., 2016). Cortical regions dmPFC, vmPFC, S1, and M1 were derived from the Harvard–Oxford atlas.

**Appendix 5—table 1.** ROI anatomical grouping, acronym, putative function in breathing regulation and interoception, and cortical lamination type (Kleckner et al., 2017).

| No. | Region | ROI | Function | Lamination |
|---|---|---|---|---|
| 1 | Subcortical | BLA | Threat & affective processing | — |
| 2 | | Piriform | Olfactory & COVID outcomes | — |
| 3 | | dPAG | Threat response, breathing drive | — |
| 4 | | vlPAG | Breathing modulation, analgesia | — |
| 5 | | Hyp | Autonomic & interoceptive control | — |
| 6 | ACC | sgACC | Visceromotor control | Agranular |
| 7 | | pACC | Visceromotor control | Agranular |
| 8 | | dACC (aMCC) | Visceromotor control | Agranular |
| 9 | Insula | dpIns | Primary interoceptive cortex | Granular |
| 10 | | dmIns | Primary interoceptive cortex | Dysgranular |
| 11 | | mvaIns | Visceromotor control | Agranular |
| 12 | | lvaIns | Sensory integration | Agranular |
| 13 | | PoI1 | Primary interoceptive cortex | Granular |
| 14 | | PoI2 | Primary interoceptive cortex | Granular |
| 15 | PFC | dmPFC | Cognitive interoceptive regulation | Granular |
| 16 | | vmPFC | Visceromotor & value processing | Agranular |
| 17 | | S1 | Somatosensory & body mapping | Granular |
| 18 | | M1 | Motor & respiratory control | Granular |

### Appendix 6 — ROI Quality Control

#### *Visual inspection*

Each mask underwent visual inspection to confirm alignment with the MNI brain template and accurate placement within subject-specific functional EPI space. Positioning of all masks was consistent with established literature, with the exception of the dpIns. The dpIns mask (Kleckner et al., 2017) is located more dorsally and posteriorly than conventional insula atlas masks, potentially corresponding to the planum temporale and including some white matter. To ensure full posterior insula coverage while maintaining consistency with prior literature, two additional posterior insula masks from the Glasser atlas (Glasser et al., 2016) were included alongside the dpIns mask (PoI1 and PoI2).

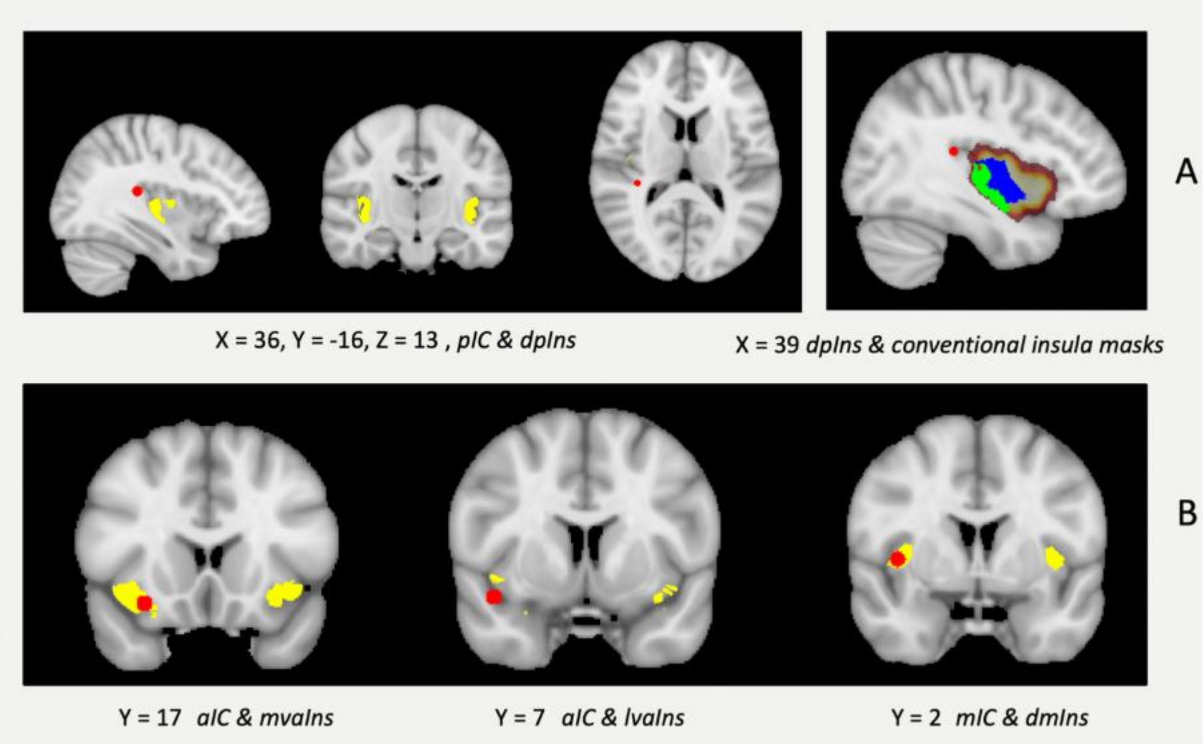


**Appendix 6—figure 1.** Seed choice of insula regions based on atlas and coordinate-derived spheres. (A) The dpIns seed lies outside the conventional cortical insula boundary. PoI1 and PoI2 were added to ensure posterior insula coverage. The ventral-posterior subdivision of the posterior insula is PoI1 in colour green. The dorsal-anterior subdivision of the posterior insula is PoI2 in colour blue. (B) The mvaIns and lvaIns seeds in colour red fall within or near the anterior insula; the dmIns seed falls within the middle insula.

***Mask volume***

Mean mask volume, voxel count, and standard deviation across subjects are reported in the table below. All ROIs exceeded 100 $mm^3$ with the exception of vmPFC in one participant (subject007), attributable to signal loss in the ventromedial region.

**Appendix 6—table 1.** Mean volume, standard deviation, and voxel number for each ROI mask in functional EPI space (1.5-mm isotropic).

| **ROI** | **Volume ($mm^3$)** | **SD** | **Voxels** |
|---|---|---|---|
| vlPAG | 311 | 39 | 92 |
| dPAG | 367 | 39 | 109 |
| Piri | 244 | 59 | 72 |
| Hyp | 845 | 222 | 250 |
| BLA | 230 | 57 | 68 |
| sgACC | 228 | 42 | 67 |
| pACC | 230 | 63 | 68 |
| dACC | 217 | 61 | 64 |
| mvaIns | 241 | 50 | 71 |
| lvaIns | 262 | 62 | 77 |
| dmIns | 229 | 40 | 68 |
| PoI1 | 4422 | 593 | 1310 |
| PoI2 | 6584 | 859 | 1951 |
| dpIns | 198 | 42 | 59 |
| dmPFC | 17572 | 4459 | 5207 |
| vmPFC | 1327 | 829 | 393 |
| S1 | 24878 | 4336 | 7371 |
| M1 | 27298 | 4192 | 8088 |

### Appendix 7 — Cohort and Site Characteristics

Because recruitment site was partially confounded with clinical subgroup, we characterised the three sub-cohorts (Oxford hospitalised, Oxford non-hospitalised, Cardiff non-hospitalised) across demographic, clinical, motion, and connectivity variables before interpreting the primary associations (Appendix 7—Table 1). Group comparisons used Kruskal–Wallis tests for continuous variables and chi-square tests for categorical variables. The sub-cohorts differed on age, sex, days since infection, hospitalisation and ventilation status (by construction), and absolute head motion. Crucially, the two breathlessness outcomes did not differ across sub-cohorts (BCS $H = 4.34$, $p = .114$; D-12 $H = 3.04$, $p = .219$), indicating comparable breathlessness burden despite clinical and demographic differences. Generalised anxiety scores (GAD-7) differed modestly across groups ($H = 6.51$, $p = .039$), being somewhat higher in the Cardiff non-hospitalised sample; this is consistent with the right-skewed GAD-7 distribution described below and is examined further in the moderation analyses. Connectivity in the dPAG–PoI1 edge did not differ across groups ($H = 0.17$, $p = .920$), whereas BLA–dACC connectivity did ($H = 7.60$, $p = .022$), motivating the site- and cohort-adjusted sensitivity models reported in the main text (Table 1).

GAD-7 scores spanned the full range of the scale (0–21; mean 6.42 ± 6.24, median 4.0) but were right-skewed (Fisher–Pearson skewness +1.15): 27 participants (51%) scored in the minimal range (<5), 14 (26%) mild (5–9), 3 (6%) moderate (10–14), and 9 (17%) severe (≥15). The higher-anxiety observations therefore clustered in a small severe subgroup rather than spreading evenly across the moderate range.

**Appendix 7—table 1.** Cohort and site characteristics by sub-cohort. Continuous imaging and demographic variables are summarised as z-scored mean ± SD. Raw symptom and anxiety scores are reported as median [IQR], categorical variables as n Group comparisons use Kruskal–Wallis H for continuous/ordinal variables and $\chi^2$ for categorical variables. Superscripts: [a] days-since-infection valid n shown in brackets; [b] FEV1/FVC valid n shown in brackets.

| Variable | Oxford Hosp. | Oxford Non-Hosp. | Cardiff Non-Hosp. | Total | Statistic | p |
|---|---|---|---|---|---|---|
| N | 19 | 9 | 25 | 53 | | |
| Age (z), mean ± SD | 0.68 ± 1.01 | 0.04 ± 0.50 | −0.53 ± 0.82 | 0.00 ± 1.00 | H=19.49 | <.001 |
| female sex, n (%) | 4 (21%) | 7 (78%) | 20 (80%) | 31 (58%) | $\chi^2$=17.11 | <.001 |
| BMI (z), mean ± SD | −0.06 ± 1.01 | 0.10 ± 0.72 | 0.01 ± 1.10 | 0.00 ± 1.00 | H=0.83 | .659 |
| Days infection-to-scan (z)[a] | −0.55 ± 0.43 [18] | −0.19 ± 0.67 [8] | 0.45 ± 1.17 [25] | −0.00 ± 1.00 [51] | H=11.02 | .004 |
| Hospitalised, n (%) | 19 (100%) | 0 (0%) | 0 (0%) | 19 (36%) | $\chi^2$=53.00 | <.001 |
| Ventilated, n (%) | 14 (74%) | 0 (0%) | 0 (0%) | 14 (26%) | $\chi^2$=34.05 | <.001 |
| BCS (raw), median [IQR] | 5.0 [1.0–13.5] | 4.0 [1.0–8.0] | 11.0 [5.0–21.0] | 8.0 [2.0–15.0] | H=4.34 | .114 |
| D-12 (raw), median [IQR] | 4.0 [1.0–10.5] | 3.0 [2.0–20.0] | 12.0 [2.0–22.0] | 7.0 [1.0–16.0] | H=3.04 | .219 |
| GAD-7 (raw), median [IQR] | 2.0 [1.5–6.5] | 3.0 [2.0–4.0] | 6.0 [4.0–16.0] | 4.0 [2.0–9.0] | H=6.51 | .039 |
| Mean FD abs. (z), mean ± SD | 0.43 ± 1.26 | −0.23 ± 0.60 | −0.24 ± 0.80 | −0.00 ± 1.00 | H=6.10 | .047 |
| Mean FD rel. (z), mean ± SD | 0.39 ± 1.16 | −0.04 ± 0.82 | −0.28 ± 0.85 | 0.00 ± 1.00 | H=4.27 | .118 |
| FEV1/FVC (z)[b] | −0.09 ± 0.28 [11] | −1.97 [1] | 0.14 ± 1.16 [21] | −0.00 ± 1.00 [33] | H=3.21 | .201 |
| dPAG–PoI1 FC, mean ± SD | −0.00 ± 0.08 | −0.00 ± 0.09 | −0.01 ± 0.11 | −0.01 ± 0.10 | H=0.17 | .920 |
| BLA–dACC FC, mean ± SD | −0.07 ± 0.10 | −0.09 ± 0.05 | −0.02 ± 0.07 | −0.05 ± 0.09 | H=7.60 | .022 |

*[a] Days infection-to-scan missing for 2 participants (valid n = 51). [b] FEV1/FVC available for 33 participants; the Oxford non-hospitalised group contributed a single valid value, so its SD*

*is undefined.*

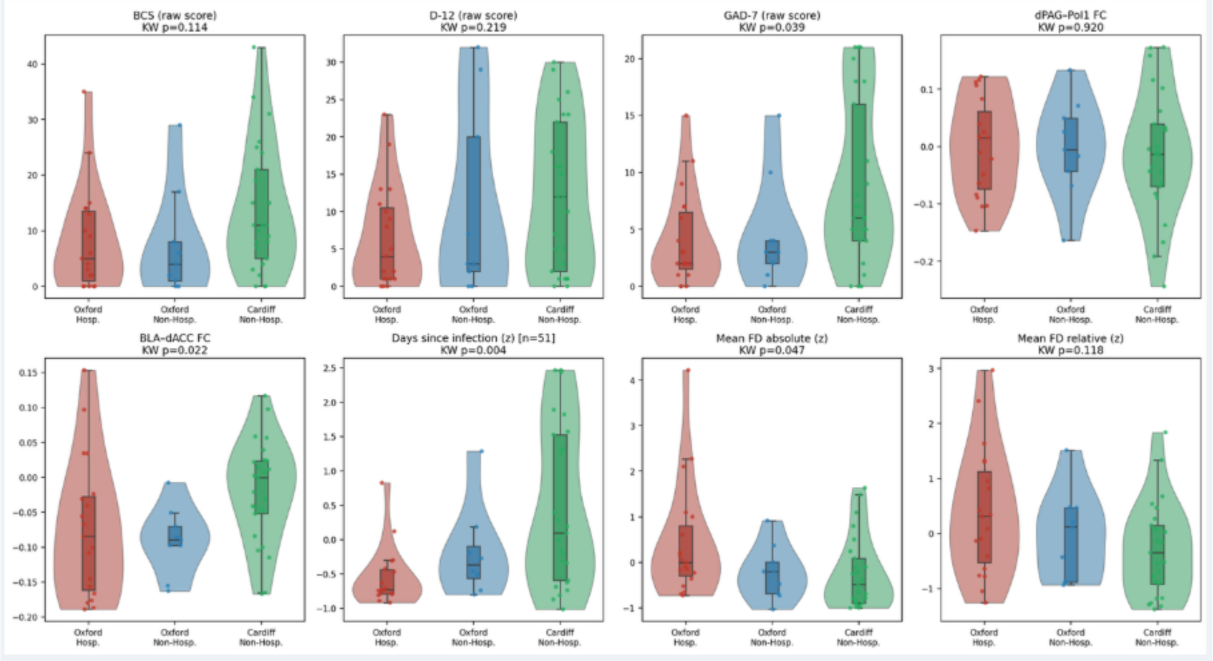


**Appendix 7—Figure 1.** Distributions of BCS, D-12, GAD-7, the two BCS-associated connectivity edges (dPAG–PoI1, BLA–dACC), days since infection, and head motion (mean absolute and relative framewise displacement) across the three sub-cohorts (Oxford hospitalised, Oxford non-hospitalised, Cardiff non-hospitalised). Each panel shows violin (distribution), box (median and interquartile range), and strip (individual participants) plots; the Kruskal–Wallis p-value is annotated above each panel. Days since infection was available for 51 of 53 participants. The two breathlessness outcomes (BCS and D-12) did not differ significantly across sub-cohorts, whereas GAD-7, BLA–dACC connectivity, days since infection, and absolute head motion did differ (all annotated per panel).

**Appendix 8 — Covariate Rationale**

The primary model was kept parsimonious because the sample size (N = 53) was modest relative to the number of candidate demographic, clinical, and imaging covariates. Inclusion decisions rested on theoretical relevance, overadjustment risk, collinearity with recruitment structure, and missing data, and not on the magnitude or significance of univariate associations with the outcomes. Accordingly, the univariate correlations below are provided descriptively, to characterise the dataset and guide sensitivity analyses, and were not used to justify excluding any covariate; a variable can confound an edge–outcome relationship even when its marginal association with the outcome is null. Continuous variables were related to outcomes using Spearman correlations ($r_s$) and binary variables using point-biserial correlations ($r^p$).

Two decisions warrant emphasis. First, generalised anxiety scores (GAD-7) showed the strongest univariate association with BCS of any non-imaging covariate ($r_s$ = .364, p = .007) yet were uncorrelated with either connectivity edge ($r_s$ = .033 and .038), the pattern expected of a downstream mediator rather than a confounder; GAD-7 was therefore reserved for the moderation analysis rather than entered as a base covariate, to avoid overadjustment. Second, variables with substantial missingness (FEV1/FVC, 38%; CRP, 25%; D-dimer, 30%; pre-existing respiratory and cardiac conditions, 23%) were excluded from the base model to avoid reducing the analytic sample and introducing selection bias, with the caveat that absence of a significant univariate association is not evidence of no effect.

Among these, pre-existing respiratory conditions merit particular attention as a plausible confound. Respiratory history was associated with both breathlessness outcomes ($r^p$ = .38, p = .01; $r^p$ = .34, p = .03) but was uncorrelated with either connectivity edge ($r^p$ = −.06, p = .73; $r^p$ = −.07, p = .66), indicating that it predicts breathlessness through a pathway independent of the functional connectivity associations reported here. This was confirmed in a dedicated sensitivity model (Table 1): adding respiratory condition as a covariate in the subsample with available data (n = 41) left both edge coefficients essentially unchanged. A more conservative analysis restricted to participants with no respiratory history (n = 31) retained the direction and approximate magnitude of both associations (dPAG–PoI1 β = −2.71, p = .119; BLA–dACC β

= 3.37, p = .101), although neither reached significance at this reduced sample size. We interpret this as a loss of power rather than evidence that the associations depend on respiratory comorbidity. Missingness in medical history was confined to the Oxford site and was unrelated to BCS, D-12, GAD-7, or either connectivity edge (all p > .10); participants with missing data were younger (p = .017), and age is a covariate in all models.

**Appendix 8—table 1.** Candidate covariates: missingness, inclusion decisions, and univariate associations with each outcome. Associations are Spearman $r_s$ for continuous covariates and point-biserial $r^p$ for binary covariates (rows marked $^p$). "Base" = inclusion in the primary model; "Sens." = evaluation in a sensitivity model; "Excl." = not entered in any model.

| Variable | In base / Sens. | Missing | r (BCS), p | r (D-12), p | r (dPAG–PoI1), p | r (BLA–dACC), p | Rationale (abbreviated) |
|---|---|---|---|---|---|---|---|
| Age | Base | 0 | −0.15, .28 | −0.12, .41 | +0.13, .37 | −0.25, .07 | Standard demographic covariate; a priori given links to lung function and FC. |
| Sex$^p$ | Base | 0 | +0.10, .46 | +0.22, .12 | +0.06, .69 | +0.09, .51 | A priori; sex differences in breathlessness perception and FC. |
| BMI | Base | 0 | +0.24, .08 | +0.05, .74 | +0.04, .79 | +0.04, .76 | A priori; BMI alters cardiorespiratory load and FC. |
| Mean FD (abs.) | Base | 0 | −0.13, .37 | −0.19, .18 | +0.20, .16 | −0.11, .43 | Standard nuisance regressor; motion inflates FC regardless of behaviour. |
| Mean FD (rel.) | Base | 0 | +0.13, .36 | −0.00, .99 | +0.19, .17 | −0.09, .51 | Complementary motion metric; dual-metric practice. |
| Site$^p$ | Sens. | 0 | +0.27, .05 | +0.19, .16 | −0.06, .66 | +0.35, .01 | Scanner/protocol variance; tested in sensitivity to avoid absorbing breathlessness variance collinear with site. |
| Cohort group$^p$ | Sens. | 0 | −0.17, .22 | −0.22, .11 | +0.05, .74 | −0.20, .16 | Collinear with hospitalisation/site; alternative deconfounding covariate. |
| Hospitalisation | Sens. | 0 | −0.17, .22 | −0.22, .11 | +0.05, .74 | −0.20, .16 | Plausibly on causal pathway; base-model exclusion avoids overadjustment for a mediator. |
| Ventilation$^p$ | Excl. | 0 | −0.01, .93 | −0.08, .55 | +0.02, .88 | −0.08, .57 | Subset of hospitalised (n = 14); near-zero associations and collinear with hospitalisation. |
| Time since infection | Sens. | 2 (4%) | −0.08, .56 | −0.21, .14 | −0.15, .30 | −0.14, .34 | Could modulate symptoms/FC; dedicated sensitivity model; 4% missing. |

| | | | | | | | |
|---|---|---|---|---|---|---|---|
| Pre-existing respiratory cond. p | Sens. | 12 (23 %) | +0.38, .01 | +0.34 , .03 | −0.06 , .73 | −0.07 , .66 | Plausible confound; associated with outcomes but not with either edge; 23% missing (one site); dedicated sensitivity model (Table 1) preserved both associations. |
| Pre-existing cardiac cond. p | Excl. | 12 (23 %) | −0.06, .72 | −0.14 , .37 | +0.08 , .62 | −0.14 , .38 | Possible contributor; 23% missing; weak associations. |
| FEV1/FVC ratio | Excl. | 20 (38 %) | −0.01, .97 | +0.16 , .38 | +0.14 , .45 | −0.03 , .88 | Objective lung pathology; 38% missing risks selection bias; weak associations. |
| CRP | Excl. | 13 (25 %) | −0.15, .36 | −0.19 , .25 | −0.04 , .80 | −0.26 , .11 | Inflammation marker; 25% missing; uncertain timing relative to scan. |
| D-dimer | Excl. | 16 (30 %) | −0.10, .58 | −0.24 , .16 | +0.05 , .79 | −0.26 , .12 | Coagulation marker; 30% missing; variable timing. |
| GAD-7 | Excl. | 0 | +0.36, .007 | +0.29 , .04 | +0.03 , .81 | +0.04 , .79 | Strongest non-brain correlate of BCS, but uncorrelated with FC; likely mediator — excluded to avoid overadjustment; used as moderator instead. |

***Note.*** *Associations are reported for descriptive and design-justification purposes only and were not used as inclusion or exclusion criteria. The hospitalisation and cohort-group rows share identical association values because, in this sample, cohort group is defined by hospitalisation status.*
